# Suppression of progesterone by influenza A virus mediates adverse maternal and fetal outcomes in mice

**DOI:** 10.1101/2023.09.11.557146

**Authors:** Patrick S. Creisher, Maclaine A. Parish, Jun Lei, Jin Liu, Jamie L. Perry, Ariana D. Campbell, Morgan L. Sherer, Irina Burd, Sabra L. Klein

## Abstract

Influenza A virus infection during pregnancy can cause adverse maternal and fetal outcomes, but the mechanism responsible remains elusive. Infection of outbred mice with 2009 H1N1 at embryonic day (E) 10 resulted in significant maternal morbidity, placental tissue damage and inflammation, fetal growth restriction, and developmental delays that lasted through weaning. Restriction of pulmonary virus replication was not inhibited during pregnancy, but infected dams had suppressed circulating and placental progesterone (P4) concentrations that were caused by H1N1-induced upregulation of pulmonary cyclooxygenase (COX)-1, but not COX-2-, dependent synthesis and secretion of prostaglandin (PG) F2α. Treatment with 17-α-hydroxyprogesterone caproate (17-OHPC), a synthetic progestin that is safe to use in pregnancy, ameliorated the adverse maternal and fetal outcomes from H1N1 infection and prevented placental cell death and inflammation. These findings highlight the therapeutic potential of progestin treatments for influenza during pregnancy.

**Importance:** Pregnant individuals are at risk of severe outcomes from both seasonal and pandemic influenza A viruses. Influenza infection during pregnancy is associated with adverse fetal outcomes at birth and adverse consequences for offspring into adulthood. We developed an outbred mouse model of 2009 H1N1 influenza virus infection during pregnancy, with semi-allogenic fetuses. When dams are infected with 2009 H1N1, in addition to pulmonary virus replication, tissue damage, and inflammation, the placenta shows evidence of transient damage and inflammation that is mediated by increased activity along the arachidonic acid pathway leading to suppression of circulating progesterone. Placental damage and suppressed progesterone are associated with long-term effects on perinatal growth and developmental delays in offspring. Treatment of H1N1-infected pregnant mice with 17-OHPC, a synthetic progestin treatment safe that is safe to use in pregnancy, prevents placental damage and inflammation and adverse fetal outcomes. This provided a novel therapeutic option for treatment of influenza during pregnancy that should be explored clinically.

## 1 Introduction

Pregnant individuals are at increased risk of severe disease and death following infection with seasonal and pandemic influenza viruses (1–5). For example, pregnancy was associated with an 18.1% increase in one hospital’s fatality rate during the 1918 H1N1 pandemic and a disproportionally greater percentage of morbidity and death during the 2009 H1N1 pandemic (1, 4). Further, seasonal influenza viruses are reported to annually infect between 483 and 1097 per 10,000 pregnant women (6), with greater odds of adverse outcomes, including intensive care unit (ICU) admission, acute cardiopulmonary morbidity, and increased hospital length of stay, among pregnant than nonpregnant people (2, 5, 7). In the absence of vertical transmission, influenza virus infection during pregnancy is still associated with adverse perinatal and fetal outcomes (8–10), as was seen during the 2009 H1N1 pandemic, where maternal influenza A virus (IAV) infection increased the relative risk of low birth weights and stillbirths by 71% and 136%, respectively (9). Moreover, infection with influenza viruses during pregnancy has been associated with long-lasting consequences for offspring, such as increasing the risk of development of neuropsychiatric and metabolic disorders in adulthood (11–13).

The 2009 H1N1 pandemic spurred development of mouse models of IAV infection during pregnancy to uncover mechanisms of adverse maternal and fetal outcomes. Collectively, these studies have revealed pregnancy-specific mechanisms of influenza pathogenesis. Notably, pregnant mice infected with diverse influenza viruses consistently experience greater morbidity and/or mortality than nonpregnant female mice, consistent with observations in humans (14–19). Pregnancy impacts pulmonary innate immune activity and tissue repair following IAV infection, as pregnant dams have greater proinflammatory cytokine concentrations and numbers of neutrophils and macrophages in lungs than nonpregnant females (16–20). Moreover, IAV infection during pregnancy is associated with pulmonary tissue damage, impaired epithelial regeneration, reduced progesterone (P4) concentrations, and a shift of alveolar macrophages from a classical to an alternative activation state (16, 20–22). Pregnant dams infected with IAV have reduced adaptive immune responses including reduced IAV-specific antibody titers, numbers of circulating CD4^+^ and CD8^+^ T cells, and IAV-specific cytotoxic T cell activity, with these effects associated with changes in circulating P4 and estrogens (17, 18, 21, 23). Whether IAV-induced changes in circulating concentrations of sex steroids are a cause or consequence of maternal disease during infection has not been determined.

Although mice do not recapitulate all aspects of human pregnancy, mice do exhibit many of the cardiovascular, hormonal, and immunological changes associated with pregnancy (24–28). For example, concentrations of P4 increase dramatically during pregnancy and support the growth of the placenta in both mice and humans (27, 29), with secretion exclusively by the corpus luteum in mice and secretion shifting from the corpus luteum to the placenta mid-gestation in humans (27, 30). Mouse models of pregnancy have limitations, including shorter gestation in mice and distinct mechanisms of placentation (25–27). Furthermore, when using inbred mice, the absence of a semi-allogeneic fetus limits tolerogenic immune mechanisms observed in humans and outbred animals (31, 32), and the presence of a semi-allogeneic fetus has been shown to increase morbidity and mortality following IAV infection (17, 33).

In the current study, using outbred mice, IAV infection during pregnancy caused significant maternal morbidity and adverse fetal outcomes that were associated with transient placental tissue damage and inflammation. Both maternal disease and adverse perinatal outcomes were mediated by IAV-induced activation of cyclooxygenase-1 (COX-1), but not COX-2, causing increased activation of prostaglandin F2α (PGF2α) and suppression of P4. Treatment of IAV-infected pregnant female mice with a synthetic progestin that is safe for administration during pregnancy reversed the adverse maternal, placental, and fetal outcomes from IAV infection. This novel study highlights utilizing hormone modulating treatments for the treatment of influenza during pregnancy.

## 2 Materials and Methods

### 2.1 Viruses and cells

Viral seed stocks of mouse adapted A/California/04/09 H1N1 (maH1N1), generated by reverse genetics from a published sequence (34) were kindly provided by Dr. Andy Pekosz (Johns Hopkins University). Working stocks were generated by infecting Madin-Darby canine kidney (MDCK) cells as described (35) and stored in aliquots at –70°C.

### 2.2 Experimental mice

Timed-pregnant adult (8–12 weeks of age) CD-1 IGS mice were purchased from Charles River Laboratories. Animals arrived at embryonic day 8 (E8) and were singly housed under standard biosafety level 2 housing conditions with ad libitum food and water. Pregnant mice were acclimated for at least 24 h prior to infections (35). Monitoring and procedures for animal experiments were performed consistently at the same time of day.

### 2.3 IAV infections and monitoring

Pregnant dams were intranasally infected at E10 with 10^3^ TCID_50_ units of maH1N1 in 30 μl of media (DMEM; Sigma D5796), a sublethal dose in nonpregnant CD-1 female mice, or mock inoculated with 30 μl of media. Prior to intranasal infection, mice were anesthetized via intraperitoneal ketamine/xylazine cocktail (80 mg/kg ketamine, 5 mg/kg xylazine). Following intranasal infections, body mass and rectal temperature were monitored once daily in the morning for 14 days or until tissue collection.

### 2.4 COX inhibitor treatments

Experimental animals were administered vehicle alone [20% DMSO (Sigma D2650) v/v diluted in PBS], the COX-1 specific inhibitor SC-560 (10 mg/kg; abcam ab120649), or the COX-2 specific inhibitor NS-398 (10 mg/kg; MedChemExpress HY-13913) in 100 μl intraperitoneally (36–38). Mice were administered treatment twice daily, starting 1 day prior to infection and until tissue collection at 3 dpi. Serum thromboxane B2 was measured to confirm COX-1 inhibition (39, 40) by ELISA per the manufacturer’s instructions (Cayman Chemical 19030). COX-2 specific enzymatic activity was measured using the cyclooxygenase activity assay kit (abcam ab204699) per the manufacturer’s instructions. Inhibitors were reconstituted and diluted fresh for each dose.

### 2.5 17α-hydroxyprogesterone caproate treatment

17α-hydroxyprogesterone caproate (17-OHPC, 250 mg/mL) was obtained from the Johns Hopkins Hospital Pharmacy (NDC 71225-105-01) and diluted to 20 mg/mL in vehicle [46% v/v benzyl benzoate (Sigma B6630) in castor oil (Sigma C9606)]. Experimental animals were administered vehicle alone or 17-OHPC subcutaneously at the time of intranasal infection. 17-OHPC was administered at a dose of 2 mg/dam, which previously demonstrated efficacy in preventing adverse neurological outcomes in a mouse model of intrauterine inflammation (41), and was administered as a single injection due to the prolonged half-life of 17-OHPC when delivered in an oil-based solution [>7 days (42)].

### 2.6 Offspring measurements and developmental assessments

Offspring from IAV- and mock-inoculated dams were measured at post-natal day (PND) 0, within 12 hours of birth. Body mass (g), length measured from nose to anus (mm), and head diameter measured from ear to ear (mm) were recorded for each pup and the average for each independent litter was calculated to avoid confounding litter effects. After birth and until weaning, pups were monitored daily for body mass (g) and the appearance of key developmental milestones including fur development, the opening of the ear canals and eyes, and teeth eruption (43, 44). At PND5, pups were subjected to the developmental neurobehavioral assay of surface righting, as described (45, 46). For each test, 1–2 male and 1–2 female offspring from at least 5 independent litters were used per condition to avoid confounding litter effects.

### 2.7 Tissue and serum collection

Experimental mice were euthanized via ketamine/xylazine overdose (160 mg/kg ketamine, 10 mg/kg xylazine) followed by cardiac exsanguination at 1, 3, 6, or 8 days post-infection. Maternal lungs, spleen, and ovaries were collected and flash frozen on dry ice for homogenization. Fetuses and placenta were flash frozen in dry ice or fixed in 4% paraformaldehyde (Thermo Fisher Scientific J19943.K2) for 72 h at 4°C for immunohistochemistry. Serum was separated from blood by centrifugation at 9,600 x g for 30Lmin at 4°C.

### 2.8 Tissue homogenization and infectious virus quantification

Frozen right cranial lungs, spleen, ovaries, placentas, and fetuses were weighed and homogenized in 1:10 weight per volume of PBS using Lysing Matrix D tubes (MP Biomedicals #6913100) in a MP Fast-prep 24 5G instrument. Homogenate was clarified by centrifugation at 12,000 rpm and stored at -70°C until analysis. IAV infectious virus titers in tissue homogenate was determined by TCID_50_ assay, as described previously (22, 35, 47), and the Reed-Muench method was used to calculate the TCID_50_ titer for each sample.

### 2.9 Multiplex cytokine analysis

Cytokines and chemokines in the right cranial lung, spleen, and placenta homogenate were multiplexed using the Immune Monitoring 48-Plex Mouse Procartaplex Panel (Thermo Fisher Scientific EPX480-20834-901) miniaturized with the Curiox DA-Bead DropArray platform (48) (**Text S1**).

### 2.10 Placental histology, immunohistochemistry, and TUNEL staining

Placentas were fixed, processed, cut, and mounted as described (46, 49). Routine H&E staining was performed to evaluate the morphological change of the placentas. Immunohistochemical staining (**Table S1**) was performed as described (46, 49). Images were taken using a Zeiss Axioplan 2 microscope (Jena) under 5x or 20x magnification. Cell density of vimentin and cytokeratin positive cell quantification was performed using Image J (1.47v) as described (46, 49). Terminal deoxynucleotidyl transferase dUTP nick end labeling (TUNEL) was performed with Click-iT™ Plus TUNEL Assay (ThermoFisher C10617) per the manufacturer protocol as described (46, 49). For each placenta, 10 random images in labyrinth at the middle level (thickest) of placenta were taken, and the average percentage of TUNEL^+^ nuclei (TUNEL^+^ nuclei divided by the total number of DAPI^+^ nuclei) given for that placenta. One placenta per dam was used and 5-9 dams per group were analyzed.

### 2.11 Flow cytometry

Placentas and maternal blood were collected from a subset of experimental dams at 6 dpi for flow cytometry as described previously (50, 51) with modifications. Briefly, single cells were prepared from 3 pooled placentas per dam. Manual digestion was performed by mincing placentas into 2-4 mm pieces, followed by enzymatic digestion with collagenase D (Roche 11088882001). Cells were passed through a 70 µm cell strainer, washed, subjected to red blood cell lysis, and Fc blocking [anti-CD16/32 (BD Biosciences 554142)]. Single cells from maternal blood (1 mL) were subjected to red blood cell lysis, and Fc blocking. Single cell suspensions were split in half and stained with one of two surface marker antibody cocktails (**Table S1**), washed, and acquired on acquired on an Attune™ NxT Acoustic Focusing Cytometer (ThermoFisher Scientific) and analyzed with FlowJo (version 10.8.1). Compensation was performed with OneComp eBeads (Thermo Fisher Scientific 01-1111-41). Gating was performed as described previously [(50, 51) and **Fig. S1**].

### 2.12 Progesterone and prostaglandin ELISA

To measure P4, steroids were extracted from serum or tissue homogenate via diethyl ether extraction at a 1:5 sample to ether ratio. Extracted steroids were diluted 1:100 (serum) or 1:10 (tissue homogenate). P4 was quantified by ELISA according to the manufacturer’s instructions (Enzo Life Sciences ADI-900-11). PGF2α (Alpco 74-PGFHU-E01) and PGE2 (Alpco 74-PG2HU-E01) in serum or tissue homogenate was quantified by ELISA according to the manufacturer’s instructions.

### 2.13 Western Blot

Protein lysates from flash frozen right caudal lung lobes and right ovaries were prepared for as described (46, 49). For each sample, 20 µg of protein was subjected to sodium dodecyl sulfate-polyacrylamide gel electrophoresis (SDS-PAGE) on NuPAGE 4-12% Bis-Tris gels, blotted onto Immobilon-FL PVDF Membrane (Millipore #IPFL00010), and the membranes blocked, washed, and stained with primary and secondary antibodies (**Table S1**) as described (46, 49). Blots were imaged on an Azure 600 imager (Azure Biosystems). Individual bands were quantified using Image Studio software (LI-COR Biosciences; version 3.1.4) The signal from each band was normalized against the GAPDH signal and graphed as arbitrary units.

### 2.14 Statistical analyses

Statistical analyses were performed using GraphPad Prism v10.0.3 (GraphPad Software). D’Agostino & Pearson tests and quantile-quantile plots were used to evaluate if data were normally distributed. If data were nonnormal or if sample size was below the minimum for D’Agostino & Pearson testing (n=8/group), non-parametric tests were used. Area under the curve (AUC) of body mass change and rectal temperature curves, lung viral titers, offspring birth measures, offspring growth AUC, offspring surface righting, and P4 concentrations were analyzed by two-tailed unpaired t-test or one-way or two-way analysis of variance (ANOVA) followed by Bonferroni *post-hoc* test. Multiplex cytokine quantifications were analyzed using multiple unpaired t-tests with Holm-Šídák correction for multiple comparisons or two-way ANOVA followed by Bonferroni *post-hoc* test. Developmental milestones, IHC quantification, TUNEL quantification, flow cytometry, western blot quantification, PG concentrations, and TxB2 concentrations were analyzed using Mann-Whitney test or Kruskal-Wallis test followed by Dunn’s *post-hoc* test. Mean or median differences were considered statistically significant at *p* < 0.05.

### 2.15 Data Availability Statement

The data supporting the conclusions of this article will be made available by the authors, without undue reservation.

## 3 Results

### 3.1 Influenza A virus infection during pregnancy causes a pulmonary cytokine storm and maternal morbidity in outbred mice

Mouse models of IAV infection during pregnancy primarily use inbred mice (14, 15, 18, 19, 22) that are devoid of fetal genetic heterogeneity (17, 33). Intranasal IAV infection of pregnant outbred CD1 mice at E10, roughly corresponding to the human second trimester (52), resulted in significant morbidity, in which IAV-infected dams gained less body mass (**Fig. 1A**) and experienced hypothermia (**Fig. 1B**) relative to mock-inoculated dams. IAV-infected dams experienced greater hypothermia than nonpregnant females (**Fig. S2A**). Subsets of dams were euthanized at 3, 6, and 8 days post infection (dpi) to measure pulmonary infectious viral loads (**Fig. 1C**) and cytokine concentrations in lungs and spleens (**Fig. 1D, Table S2, Table S3**) prior to and during peak disease. Pulmonary viral load peaked at 3 dpi and was low to undetectable at 8 dpi (**Fig. 1C**), which was consistent and not significantly different from IAV-infected nonpregnant female mice (**Fig. S2B**). Prototypical cytokines and chemokines associated with influenza (53), including interferons (IFNs), tumor necrosis factor (TNF) α, interleukin (IL)-6, and CCL2 were upregulated throughout infection in the lungs, but not the spleens of IAV-infected dams (**Fig. 1D, Table S2, Table S3**). Together, these data indicate that outbred pregnant mice infected with IAV experience morbidity to a greater degree than nonpregnant females and a pulmonary cytokine storm, consistent with previous studies (14, 18, 22).

**Figure 1.**
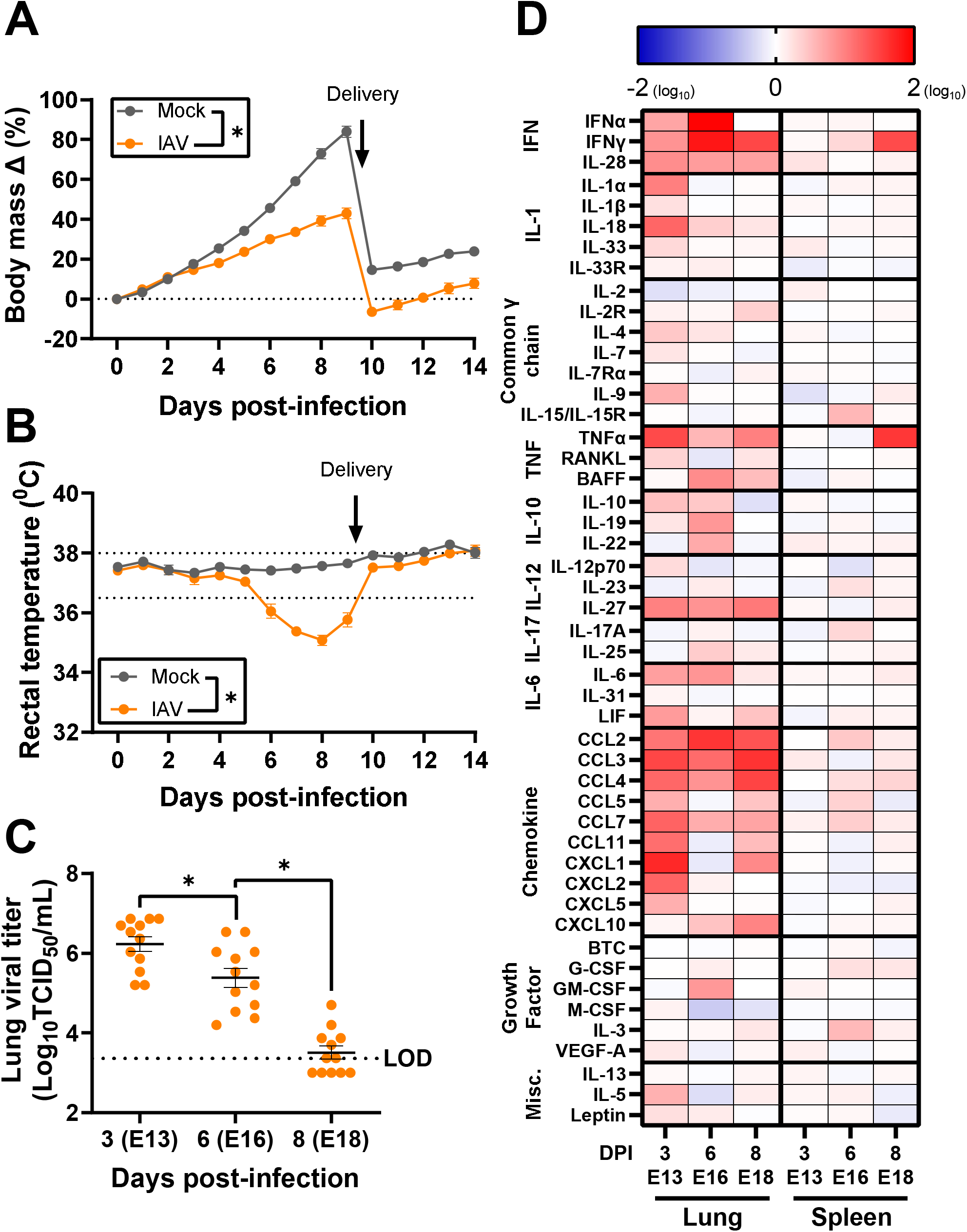
Influenza A virus infection of pregnant outbred mice results in morbidity and a pulmonary cytokine storm. Dams at embryonic day (E) 10 were intranasally inoculated with 10^3^ TCID_50_ of 2009 H1N1 or were mock inoculated with media. Daily maternal body mass (A) and rectal temperature (B) were recorded as measures of maternal morbidity for 14 days. Subsets of dams were euthanized to measure viral titers in lung tissue via TCID_50_ assay (C) and cytokine concentrations in spleen and lung tissue (D) at 3 (E13), 6 (E16), and 8 (E18) days post infection. Cytokines were multiplexed with the Immune Monitoring 48-Plex Mouse Procartaplex Panel miniaturized with the Curiox DA-Bead DropArray platform (n=12-14/group). Individual symbols (A-B) or bars (C) represent the mean ± standard error of the mean per group from two independent replications (n =12-14 mice/group) with individual mice indicated by symbols in (C). Significant differences (p < 0.05) were determined by two tailed unpaired t-test of AUCs (A-B) or one-way ANOVA with Bonferroni post hoc test (C) and are indicated by an asterisk (*).

### 3.2 Influenza A virus infection during pregnancy causes intrauterine growth restriction and postnatal developmental delays

IAV infection during pregnancy is associated with adverse fetal outcomes including pre-term delivery, fetal growth restriction, and low birthweights (8–10, 54) as well as long-term health consequences in adulthood (11, 55) in humans. In mice, IAV infection during pregnancy did not impact litter size (**Fig. 2A**) or increase the rate of stillbirths (98.78% viability) versus mock inoculated dams (98.82% viability). IAV infection of dams resulted in significant fetal growth restriction, with pups of IAV-infected dams being smaller at birth based on measures of mass, length, and head diameter, than offspring of mock-inoculated dams (**Fig. 2B-D**). Offspring of IAV-infected dams remained smaller than offspring from mock-inoculated dams through weaning (**Fig. 2E**). Offspring of IAV-infected dams also displayed developmental delays, reaching key developmental milestones (44) on average 1-2 days later than offspring of mock-inoculated dams (**Fig. 2F**). Impaired surface righting, a developmental reflex (44, 45), also was apparent, with offspring, particularly males, of IAV-infected dams taking significantly longer to right themselves (**Fig. 2G**). Fetal growth restriction and developmental delays were not caused by pre-term birth or impaired postnatal care as all dams, regardless of maternal infection, delivered pups at approximately E20 (56) and displayed appropriate pup retrieval (57, 58) at PND1 and PND7 (**Fig. S3**). These data indicate that IAV infection of outbred mice causes both short and long-term adverse perinatal and fetal outcomes, consistent with humans (9, 55) and mice (15, 19, 22, 33).

**Figure 2.**
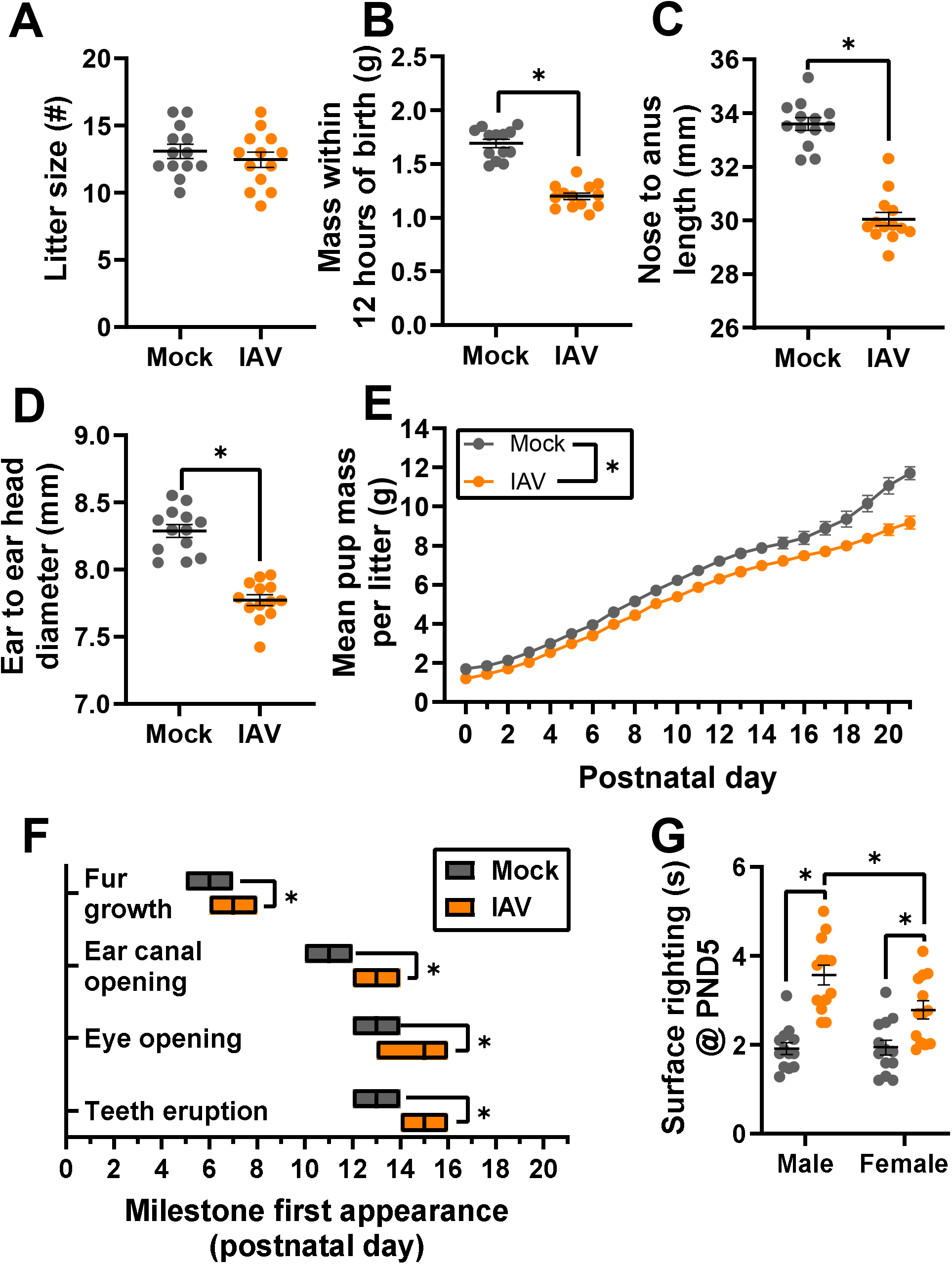
Influenza A virus (IAV) infection during pregnancy causes offspring growth restriction and developmental delays. Dams at embryonic day (E) 10 were intranasally inoculated with 10^3^ TCID_50_ of 2009 H1N1 or were mock inoculated with media and were followed through delivery. Offspring were followed through postnatal day (PND) 21 to assess fetal outcomes. Litter size (A), mass (B), nose to anus length (C), and ear to ear head diameter (D) were measured at PND 0. Average measurements of each independent litter were graphed to account for litter effects. Average pup mass for each litter was taken daily to track growth through PND 21 (E). Developmental milestones including fur growth, ear canal opening, eye opening, and teeth eruption were recorded (F). At PND 5, a subset of pups was sexed and assessed for their surface righting ability to measure neurological development. One pup per sex per dam was given 3 trials given for the test, and each pup’s best trial was reported (G). Bars (A-D, G) or symbols (E) represent the mean ± standard error of the mean per group from two independent replications (n =13 litters or pups/group) with individual litter means (A-D) or individual mice (G) indicated by symbols. For developmental milestones, the center bar of the floating boxplot indicates the median day a litter reached each milestone (n=13 litters/group), with the range indicated by the edges of the box (D). Significant differences (p < 0.05) were determined by two tailed unpaired t-test (A-D), two tailed unpaired t-test of AUCs (E), Mann-Whitney nonparametric test (F), or two-way ANOVA with Bonferroni post hoc test (G) and are indicated by an asterisk (*).

### 3.3 Influenza A virus infection during pregnancy causes transient but significant placental cell death and trophoblast loss

While many models of IAV infection during pregnancy have focused on uncovering mechanisms of maternal disease (16, 17, 19, 21, 22), the mechanisms of adverse fetal outcomes remain less clear. Placental tissue damage, in the absence of direct viral infection, has been reported in mice (15, 18, 21) and in limited observational human studies (59, 60). Despite no infectious virus in the placenta (data not shown), at 3 dpi, there was significant placental damage, including exposure of fetal red blood cells to maternal red blood cells (**Fig. 3A**), suggestive of disruption of the placental trophoblast-endothelial cell barrier, as well as increased cell death as compared with placentas from mock-inoculated dams (TUNEL^+^ cells, **Fig. 3B** for representative image, **Fig. 3E** for quantification). Staining for cytokeratin (trophoblasts, **Fig. 3C** for representative image, **Fig. F** for quantification) and vimentin (endothelial cells, **Fig. 3D** for representative image, **Fig. 3G** for quantification) revealed a loss of trophoblasts, but not endothelial cells, in placentas from IAV-infected compared with mock-inoculated dams at 3 dpi. Concentrations of IL-13 (**Fig. 3H**), granulocyte-macrophage colony-stimulating factor (GM-CSF, **Fig. 3I**), and IL-18 (**Fig. 3J**), cytokines secreted by trophoblasts *in vitro* (61, 62), were lower in placentas from IAV-infected than mock-inoculated dams at 3 dpi. By 6 and 8 dpi, neither trophoblast nor endothelial cell numbers differed between placentas of IAV-infected and mock-inoculated dams (**Fig. S4**), indicating resolution of placental damage.

**Figure 3.**
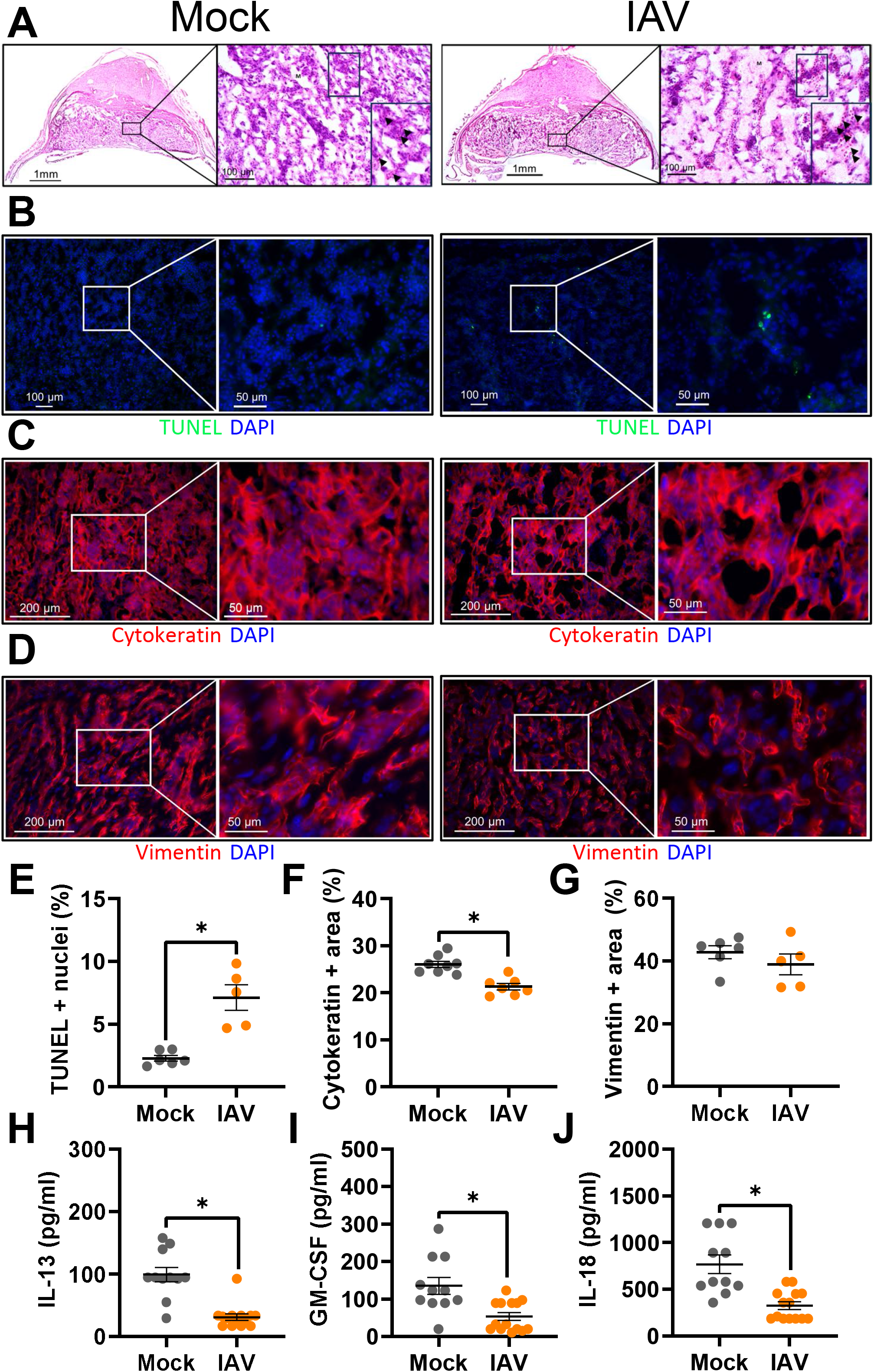
Influenza A virus infection during pregnancy causes placental damage and disruption to the trophoblast-endothelial cell barrier. Dams at embryonic day (E) 10 were intranasally inoculated with 10^3^ TCID_50_ of 2009 H1N1 or were mock inoculated with media and were euthanized at 3 dpi (E13) to collect placentas. Representative H&E images were taken at 5x magnification and 20x magnification, with the boxed portion further enlarged. Arrowhead indicate fetal red blood cells and “M” indicates maternal circulation with non-nucleated red blood cells (A). Representative TUNEL^+^ (green) stained placentas with DAPI to label nuclei taken (blue) were taken at 20x magnification and 100x (B). Placentas were immunostained for cytokeratin (C, red) to mark trophoblasts or vimentin (D, red) to mark endothelial cells and DAPI (blue) to label nuclei. Representative images were taken at 20x magnification and further zoomed 2.90-fold. Quantification of the percentage of positive nuclei (E) or positive area for each marker is shown (F-G). Placentas were homogenized and analyzed for concentrations (pg/ml) of IL-13 (H), GM-CSF (I), and IL-18 (J). Bars represent the mean ± standard error of the mean per group [n =5-6 mice/group (image quantification) and 11-14 mice/group (cytokines)] with each symbol indicating one placenta and, for analysis of images, is the mean quantification of 6 fields of view. Significant differences (*p* < 0.05) were determined by Mann-Whitney nonparametric test (E-G) or unpaired two tailed t-test (H-J) and are indicated by an asterisk (*). Scale bar: 1 mm (A, left panel/group), 100 µm [(A, right panels/group; B, left panel/group), 200 µm (C-D, left panel/group), or 50 µm (C-D, right panel/group).

### 3.4 Proinflammatory cytokines peak in the placenta following placental damage during Influenza A virus infection

Proinflammatory cytokines, such as IL-1β and TNFα, in the placenta are associated with placental damage (49, 63, 64). Concentrations of proinflammatory cytokines, including IL-1β (**Fig. 4A**), IL-6 (**Fig. 4B**), and TNFα (**Fig. 4C**) exhibited a transient increase in placentas from IAV-infected compared with mock-inoculated dams at 6dpi. The increased concentrations of proinflammatory cytokines followed the placental damage and reductions in trophoblasts observed at 3dpi. The profile of cytokine secretion in the placenta (**Table S4**) differed from the patterns observed in either the spleen (**Table S3**) or lungs (**Table S2**). Increased secretion of proinflammatory cytokines in the placenta could be mediated by infiltration of immune cells, as we have observed in models of intrauterine inflammation (50, 51). Flow cytometry was used to evaluate innate (i.e., NK cells, neutrophils, eosinophils, macrophages) and adaptive (i.e., B cells, T cells) immune cells in the placenta at 6 dpi (**Fig. 4D-L, Fig. S1** for representative gating). Frequencies of CD45^+^ leukocytes were reduced in placentas of IAV-infected dams compared to mock-inoculated dams (**Fig 4D**), which was further observed for NK cells (**Fig. 4E**), B cells (**Fig. 4F**), CD4^+^ T cells (**Fig. 4G**), and CD8^+^ T cells (**Fig. 4H**) in placentas of IAV-infected dams. Frequencies of neutrophils (**Fig. 4I**), eosinophils (**Fig. 4J**) and CD163^+^ macrophages (**Fig. 4K**) were similar in placentas from IAV- and mock-inoculated dams. CD68^+^ macrophages were the only immune cell population that exhibited a significant increase in placentas of IAV-infected compared with mock-inoculated dams (**Fig. 4L**). The patterns of immune cell infiltrates observed in placentas were not observed in peripheral blood cells obtained from the same dams (**Fig. S5**). CD68^+^ macrophages in the placenta are associated with intrauterine growth restriction in humans (65) and secretion of proinflammatory cytokines (66). Taken together, these results indicate that maternal respiratory infection with IAV can induce transient but significant cell death and proinflammatory cytokine release at the maternal fetal interface, in the absence of direct viral infection or broad leukocyte infiltration.

**Figure 4.**
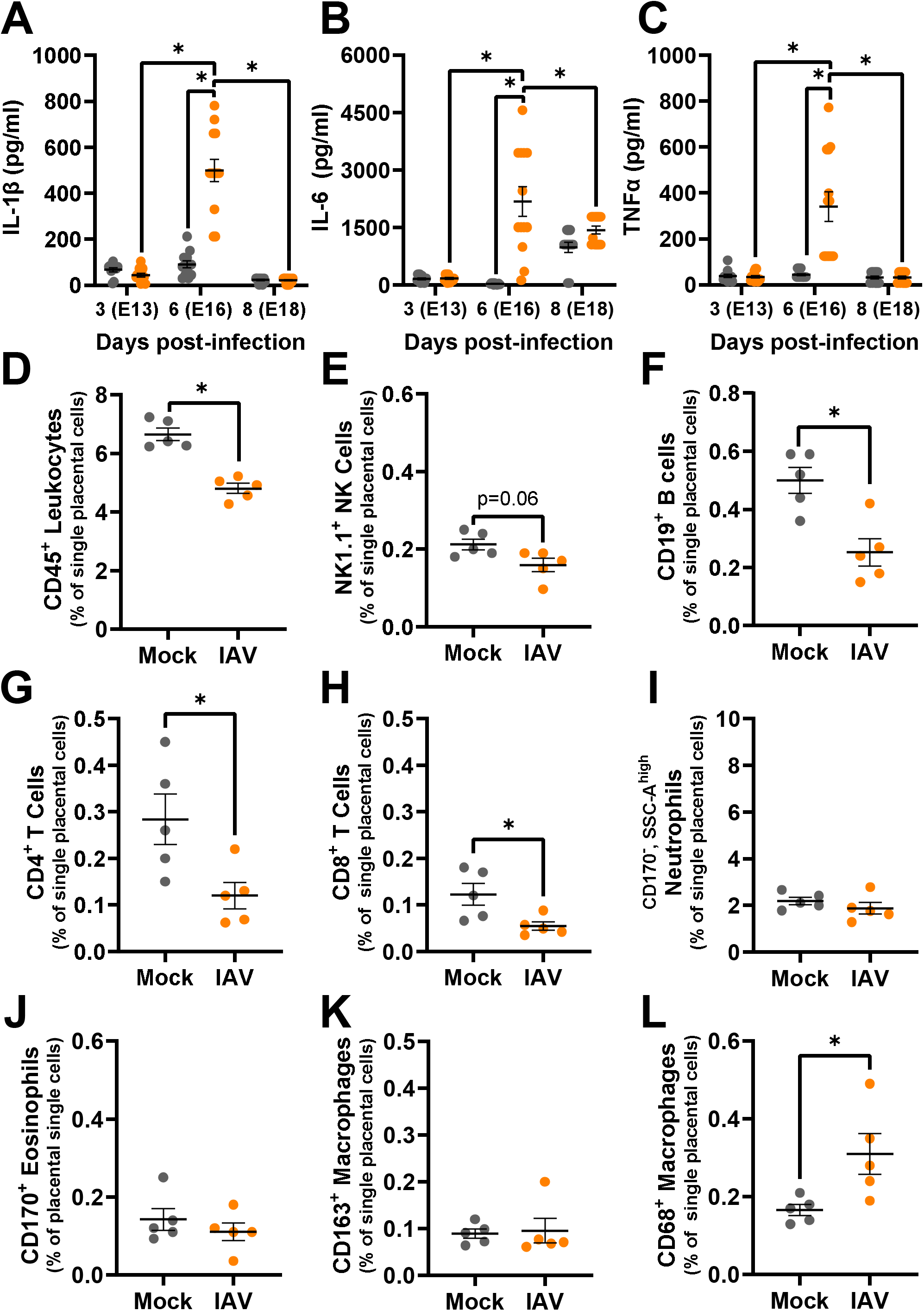
Placental concentrations of proinflammatory cytokines and immune cells peak at 6 days-post infection. Dams at embryonic day 10 were intranasally inoculated with 10^3^ TCID_50_ of 2009 H1N1 or were mock inoculated with media and were euthanized at 3 (E13), 6 (E16), or 8 dpi (E18) to collect placental tissue (A-C). Placentas were homogenized and analyzed for concentrations (pg/ml) of proinflammatory cytokines, including IL-1β (A), IL-6 (B), and TNFα (C). Flow cytometry was performed using live placental cells at 6 dpi to assess immune cell type frequencies in placental tissue including leukocytes (CD45^+^, D), NK cells (CD45^+^, CD3^-^,CD19^+^, E), B cells (CD45^+^, CD3^-^,CD19^+^, F), CD4^+^ T-cells (CD45^+^, CD3^+^, CD4^+^, G), CD8^+^ T cells (CD45^+^, CD3^+^, CD8a^+^, H), neutrophils (CD45^+^, CD11b^+^, F4/80^-^, CD170^-^, side scatter area high, I), eosinophils (CD45^+^, CD11b^+^, F4/80^+^, CD170^+^, J), CD163^+^ macrophages (CD45^+^, CD11b^+^, F4/80^+^, CD163^+^, K), and CD68+ macrophages (CD45^+^, CD11b^+^, F4/80^+^, CD68^+^, L). Bars represent the mean ± standard error of the mean per group from two independent replications (n =11-14/ mice group for cytokines and 5 mice/group for flow cytometry) with individual mice indicated by symbols. Significant differences (p < 0.05) were determined by two-way ANOVA with Bonferroni post hoc test (A-C) or Mann-Whitney nonparametric test (D-L) and are indicated by an asterisk (*).

### 3.5 Influenza A virus infection suppresses the pregnancy-associated rise in progesterone via COX-1 activity

The mechanism underlying placental damage and cytokine secretion during IAV infection might involve sex hormone regulation. Placental development and maturation, which occurs at E10-E14 in mice (67) are supported by P4 signaling in both humans and mice (27, 29, 30, 68). As IAV infection during pregnancy has been associated with reduced circulating P4 in mice (14, 15, 22), we evaluated if IAV infection of outbred pregnant mice altered circulating (**Fig. 5A**) and placental (**Fig. 5B**) P4 concentrations. Circulating and placental concentrations of P4 in mock-inoculated dams increased throughout gestation and peaked at approximately E16 (**Fig. 5A-B**) (69). In contrast, IAV infection significantly reduced circulating and placental P4 concentrations throughout gestation relative to mock-inoculated dams, although not to circulating concentrations observed in nonpregnant females **(Fig. 5A**).

**Figure 5.**
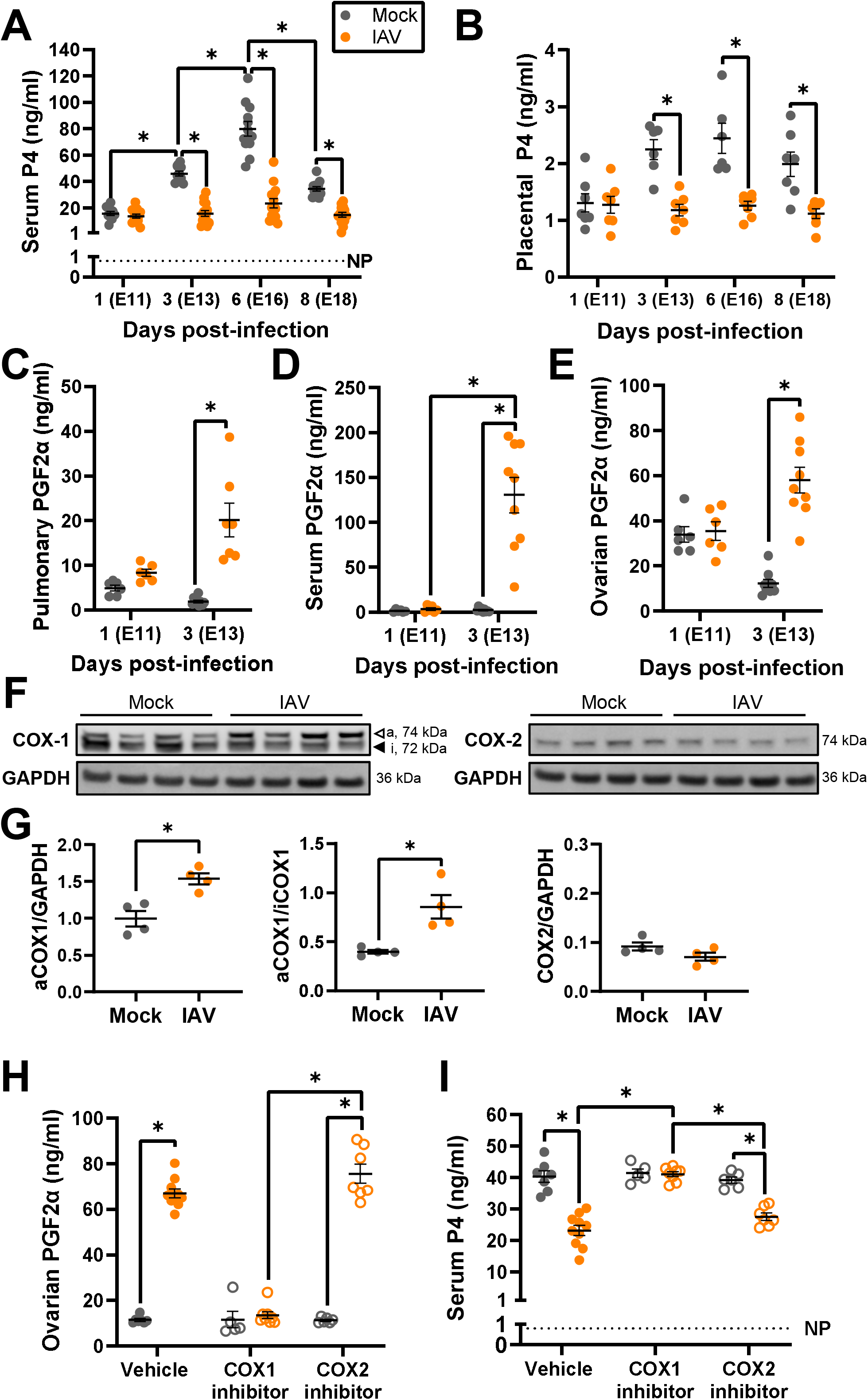
Downregulation of circulating and placental P4 caused by upregulation of COX-1 through PGF2α secretion. Dams at embryonic day (E) 10 were intranasally inoculated with 10^3^ TCID_50_ of 2009 H1N1 or were mock inoculated with media. Dams were euthanized for tissue and serum collection at 1 (E11), 3 (E13), 6 (E16) or 8 (E18) dpi. Serum (A) and placental (B) P4 concentrations (ng/ml) were measured by ELISA. PGF2α concentrations (ng/ml) were measured by ELISA in lung homogenates (C), serum (D), and ovarian homogenates (E). COX-1 and COX-2 protein expression in lung tissue was assessed by western blot at 3dpi (F) and quantified as the fluorescence signal for each protein normalized to GAPDH. Active COX-1 is also shown as a ratio to inactive COX-1 (F). Lungs from two independent experiments were analyzed together to minimize interassay variability. Independent cohorts of dams were intranasally inoculated with 10^3^ TCID_50_ of 2009 H1N1 or were mock inoculated with media and treated twice daily beginning 1 day before infection with either COX-1 inhibitor (SC-560, 10 mg/kg), COX-2 inhibitor (NS-398, 10 mg/kg), or vehicle until dams were euthanized at 3 dpi for tissue and serum collection. PGF2α concentrations (ng/ml) were measured by ELISA in ovarian tissue (H) and P4 concentrations (ng/ml) were measured by ELISA in serum (I). Bars represent the mean ± standard error of the mean per group from two independent replications (n =6-12 mice /group for ELISAs and 4 mice/group for western blot) with individual mice indicated by symbols. Significant differences (p < 0.05) were determined by two-way ANOVA with Bonferroni post hoc test (A-B, I), Kruskal–Wallis with Dunn’s post hoc test (C-E, H) or Mann-Whitney nonparametric test (G) and are indicated by an asterisk (*).

In mice, P4 is secreted by the corpus luteum of the ovary throughout the entire pregnancy where it is canonically regulated by the prostaglandin (PG) F2α (23, 70). To evaluate if secreted PGF2α upregulation was associated with reduced P4, PGF2α as well as PGE2, a prostaglandin that has been observed to be upregulated with IAV infection in nonpregnant mice (71), were measured in the lungs (i.e., the site of IAV replication), systemic circulation, and the ovaries (i.e., the site of P4 secretion) of IAV-infected and mock inoculated dams at 1 and 3 dpi (**Fig. 5C-E, Fig. S6**). Concentrations of PGF2α, but not PGE2 (**Fig. S6A-C**), were significantly increased in the lung (**Fig. 5C**), sera (**Fig. 5D**), and ovaries (**Fig. 5E**) of IAV-infected dams as compared with mock-inoculated dams at 3 dpi. Elevated concentrations of PGFα in the ovary were associated with reduced serum P4 in both IAV-infected and mock-inoculated dams (**Fig. S6D**).

Prostaglandins, including PGF2α are canonically synthesized by COX enzymes, with COX-1 constitutively expressed in most cells and COX2 induced by inflammatory stimuli (72). Active COX-1 (73), but not COX-2, expression was elevated in the lungs of IAV-infected relative to mock-inoculated dams at 3dpi (**Fig. 5F-G**, **Fig. S7**). Neither COX-1 nor COX-2 expression were altered in the ovaries (**Fig. S8**), suggesting that PGF2α generation by COX-1 is local at the site of IAV infection. To evaluate if COX-1 activity is not just sufficient but necessary for PGF2α secretion and suppression of P4 during IAV-infection, subsets of IAV-infected or mock-inoculated dams at E10 were pretreated with either a COX-1-specific inhibitor (SC560), a COX-2-specific inhibitor (NS-398), or vehicle (**Fig. 5H-I**). COX-1 inhibition, as measured by inhibition of serum thromboxane B2 (TXB2) (39, 40), was specific to COX-1 inhibitor treatment and not observed with COX-2 inhibitor treatment (**Fig. S9A**). COX-2 enzymatic activity was minimal to undetectable in the lungs of IAV infected dams regardless of treatment (**Fig. S9B**). COX-1, but not COX-2, inhibition significantly reduced PGF2α and increased P4 secretion in IAV-infected dams (**Fig. 5H-I, Fig. S9C**). These data illustrate that IAV infection disrupts activity along the arachidonic acid pathway in pregnant dams to cause suppression of P4, which is required for a healthy pregnancy.

### 3.6 Synthetic progestin treatment improves IAV-induced maternal morbidity, placental damage, and perinatal developmental delays

To establish whether IAV infection-induced P4 suppression was the cause of adverse maternal and fetal outcomes, a subset of IAV-infected and mock inoculated dams were treated with a single dose of either 17-α-hydroxyprogesterone caproate (17-OHPC), a synthetic progestin that is safe to use during pregnancy (74) or vehicle. The administration of 17-OHPC to mock-inoculated dams had no adverse impact on pregnancy outcomes. Administration of 17-OHPC to IAV-infected dams significantly reduced hypothermia (**Fig. 6A**) and increased maternal weight gain (**Fig. S10A**) as compared with IAV-infected dams treated with vehicle. Treatment of IAV-infected dams with 17-OHPC did not affect pulmonary infectious virus titers (**Fig. 6B**). These data suggest that IAV-induced suppression of P4 contributes to maternal morbidity, independent of controlling virus replication.

**Figure 6.**
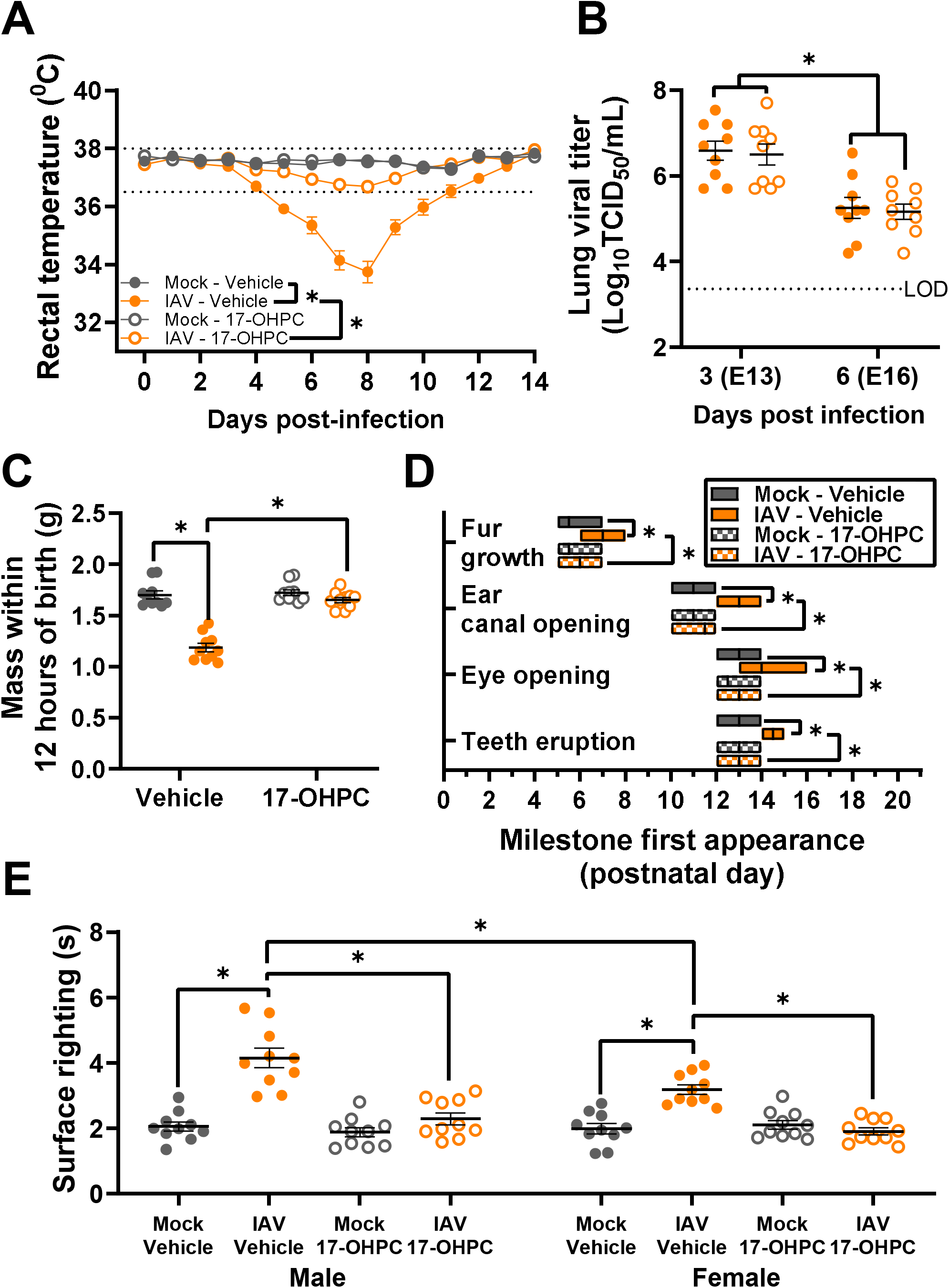
Synthetic progestin treatment improves maternal morbidity and prevents offspring growth restriction and developmental delays. Dams at embryonic day (E) 10 were intranasally inoculated with 10^3^ TCID_50_ of 2009 H1N1 or were mock inoculated with media. At the time of inoculation, dams were subcutaneously administered 17α-hydroxyprogesterone caproate (17-OHPC, 2mg/dam). Dams were euthanized for tissue collection at 3 (E13) or 6 (E16) days post infection (dpi) or carried their pregnancy through delivery (E20). Daily rectal temperatures were recorded as a measurement of maternal morbidity through 14dpi (A). Viral titers were measured in lung tissue via TCID_50_ assay at 3 (E13) and 6 (E16) dpi (B). At postnatal day (PND) 0, pups were weighed, with the average measurements of each independent litter graphed to account for litter effects (C), and pups were followed through weaning to observe developmental milestones (D) including fur growth, ear canal opening, eye opening, and teeth eruption. (E) At PND 5, a subset of pups was sexed and assessed for their surface righting ability with one pup per sex per dam given 3 trials for the test, and each pup’s best trial reported (G). Individual symbols (A) or bars (B-C, E) represent the mean ± standard error of the mean per group from two independent replications (n =10 mice or litters /group) with individual mice (B, E) or individual litter means (C) indicated by symbols. For developmental milestones, the center bar of the floating boxplot indicates the median day a littler reached each milestone (n=10 litters/group), with the range indicated by the edges of the box (D). Significant differences (p < 0.05) were determined by two-way ANOVA with Bonferroni post hoc test of AUCs (A), two-way ANOVA with Bonferroni post hoc test (B-C), Kruskal-Wallis with Dunn’s post hoc test (D), or three-way ANOVA with Bonferroni post hoc test (G) and are indicated by an asterisk (*).

To determine if administration of 17-OHPC to IAV-infected dams could reverse adverse perinatal outcomes, subsets of IAV-infected and mock-inoculated dams with and without 17-OHPC treatment carried their pregnancies to term. Treatment of mock-inoculated dams with 17-OHPC had no effect on litter size (**Fig. S10B**) or the development of offspring relative to vehicle-treated, mock-infected dams. In contrast, treatment of IAV-infected dams with 17-OHPC significantly increased birth mass (**Fig. 6C**) as well as body length and head diameter (**Fig. S10C-D**), mitigated developmental delays in key milestones (**Fig. 6D**), improved growth through weaning (**Fig. S10E**), and improved neurobehavioral development as measured by surface righting at PND5 (**Fig. 6E**) as compared with offspring from IAV-infected dams that were treated with vehicle. These data demonstrate that IAV-induced suppression of P4 during pregnancy mediates intrauterine growth restriction and postnatal developmental delays, which are reversed by treatment with a safe, synthetic progestin.

We hypothesized that 17-OHPC treatment might prevent placental damage to improve offspring outcomes among IAV-infected dams. Treatment of mock-inoculated dams with 17-OHPC did not alter placental histology, cell death, or numbers of trophoblasts as compared with vehicle-treated, mock-infected dams. In contrast, treatment of IAV-infected dams with 17-OHPC significantly reduced placental cell death (**Fig. 7A-B, Fig. S11**), increased proportions of trophoblasts (**Fig. 7C-D**), and reduced placental concentrations of proinflammatory cytokines (**Table S5**), including IL-1β (**Fig. 7E**), IL-6 (**Fig. 7F**), and TNFα (**Fig. 7G**). These data illustrate that suppressed P4 is the cause of placental tissue damage and adverse perinatal outcomes during IAV-infection and treatment with 17-OHPC improved maternal, placental, and offspring outcomes during infection.

**Figure 7.**
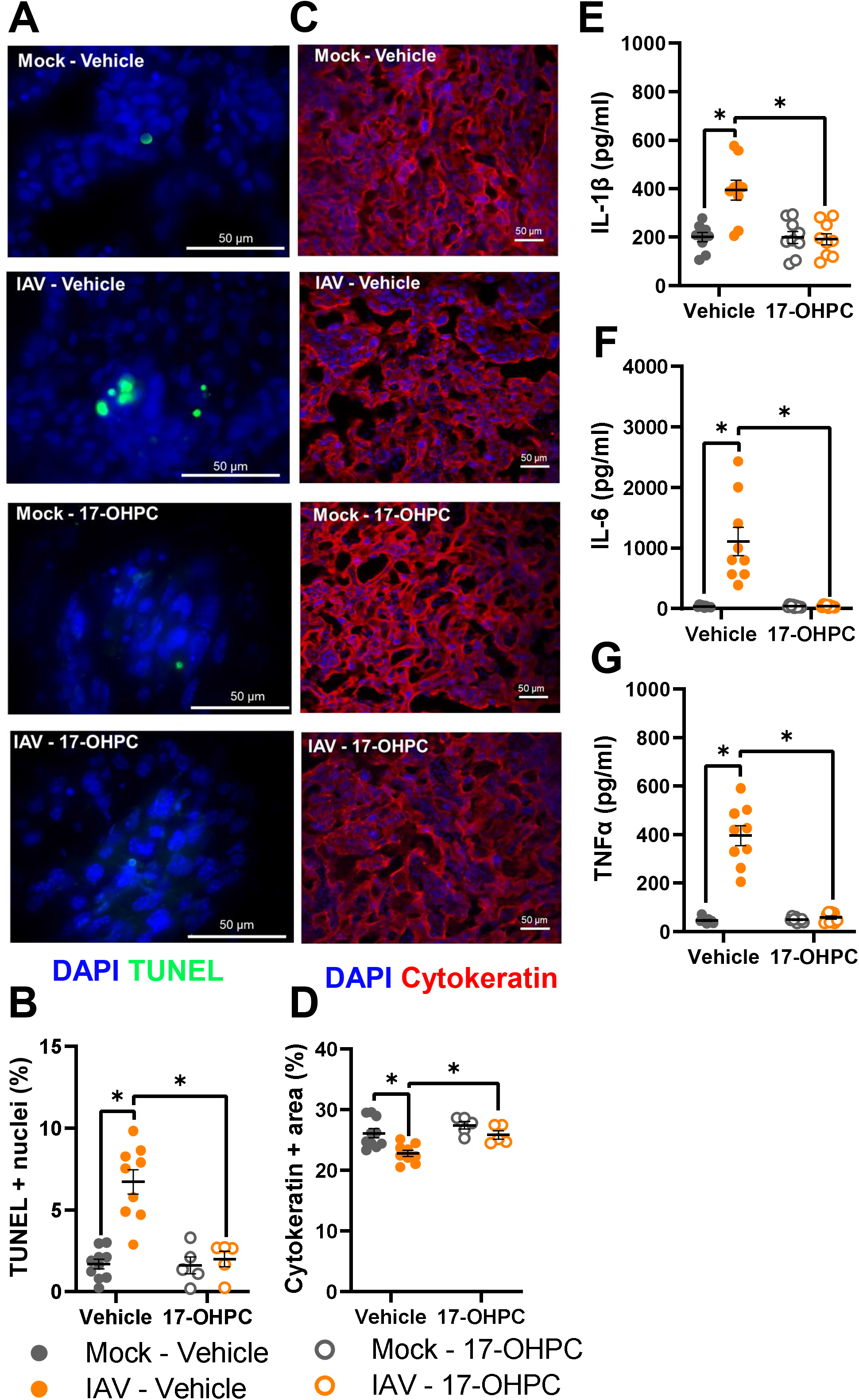
Synthetic progestin treatment ameliorates placental damage and inflammation. Dams at embryonic day (E) 10 were intranasally inoculated with 10^3^ TCID_50_ of 2009 H1N1 or were mock inoculated with media. At the time of inoculation, dams were subcutaneously administered 17α-hydroxyprogesterone caproate (17-OHPC, 2mg/dam). Placentas were collected at 3 (E13) or 6 (E16) days post infection (dpi). Representative TUNEL^+^ (green) stained placentas with DAPI to label nuclei taken (blue) at 100x magnification (A), with quantification of the percentage positive nuclei (B). Placentas were immunostained for cytokeratin (red) to mark trophoblasts and DAPI (blue) to label nuclei (C) Representative images were taken at 20x magnification, with quantification of the percentage positive area (D). Placentas were homogenized to measure concentrations (pg/ml) of IL-1β (E), IL-6 (F), and TNFα (G). Bars represent the mean ± standard error of the mean per group (n =5-9 mice /group) with each symbol indicating one placenta and, for analysis of images, is the mean quantification of 6 fields of view. Significant differences (*p* < 0.05) were determined by Kruskal-Wallis with Dunn’s post hoc test and are indicated by an asterisk (*). Scale bar: 50 µm

## 4 Discussion

Both seasonal and pandemic influenza virus infection during pregnancy result in adverse perinatal and long-term detrimental outcomes that have been reported since the 1918 pandemic (9, 55, 75). Traditionally, mouse models of IAV during pregnancy have focused on mechanisms of maternal disease (16, 19, 22) rather than characterization of the mechanisms of adverse fetal outcomes. Using an outbred mouse model of IAV infection during pregnancy, 2009 H1N1 infection caused maternal morbidity, transient placental damage and inflammation, and long-term offspring growth restriction and developmental delays. The IAV-induced adverse maternal and fetal outcomes were mediated by a COX-1-dependent reduction in P4, including at the maternal-fetal interface. Previous studies report damage to tissue architecture, necrosis, and pro-inflammatory cytokines in the placenta following maternal IAV infection (18, 76, 77), including associations between placental damage and reduced placental P4 (15, 21). Our study expands upon this literature by defining the temporal kinetics of progesterone suppression, placental cell death, and proinflammatory cytokine secretion and by establishing a novel causal role for the arachidonic acid biosynthesis pathway, which is activated by IAV infection and disrupts P4 during pregnancy. Our data provide a mechanistic connection between a pulmonary virus infection, which causes elevated COX-1 enzymatic activity, and placental tissue damage, caused by increased prostaglandin (PG) and reduced P4 synthesis. Our results are consistent with knockout studies where selective disruption of progesterone receptor signaling in immune cells causes damage to the placenta, intrauterine growth restriction, and fetal wasting (78, 79). By treating dams with a synthetic progestin that is safe during pregnancy, the IAV-induced placental damage, inflammation, and adverse fetal outcomes, including developmental delays, were reversed, highlighting the translational significance of these findings.

IAV-induced suppression of P4 was dependent on COX-1, but not COX-2, signaling and PGF2α secretion. Progesterone secretion during pregnancy is canonically regulated by PGF2α, which signals in the corpus luteum and induces luteolysis to inhibit P4 secretion (70). PGF2α is generated within the arachidonic acid biosynthesis pathway, where COX enzymes, predominantly COX-1 or COX-2, convert arachidonic acid into prostanoids that are then converted to PG subclasses (i.e., PGF2α) by specific PG synthases (80, 81). Historically, COX-1 has been characterized as constitutively active and homeostatic, while COX-2 has been characterized as inducible and associated with pain and inflammation (80, 82). COX-1, however, also may have functions in inflammatory responses, as COX-1 inhibition can alleviate neuroinflammation in a mouse model of retinitis pigmentosa (83). Further, aspirin given at COX-1-specific doses is associated with reduced rates of preeclampsia (84). Our data add to this growing body of literature by illustrating that IAV infection upregulates COX-1 in the lungs and increased COX-1-dependent PGF2α in several tissues, including the ovaries.

Mouse models of pregnancy are powerful tools to identify mechanisms of disease pathogenesis and evaluate the safety and efficacy of potential treatments (85); mice, however, have limitations. The full-term gestation for mice is approximately 20 days (about 3 weeks), but 40 weeks (about 9 months) for humans, with disproportionally more gestational time spent in the first-trimester equivalent in mice than in humans (52). Further, placental anatomy differs between mice and humans [reviewed in (28)], with, for example, the placental labyrinth structure found in mice but not humans. Inbred mice further lack genetic heterogeneity at the maternal-fetal interface, and in the current study, outbred mice were used to avoid this limitation (86). With the use of outbred mice, genetic manipulations were not available to test hypotheses; instead, pharmacologic inhibitors were used, which could have off-target effects, despite dosages optimized for target-specificity.

Despite these limitations, results from this study highlight the value of treatments that mitigate disease in addition to antiviral treatments for influenza during pregnancy. In the current study, we utilized 17-OHPC, a synthetic progestin that has been prescribed to pregnant women with history of preterm birth (87). While the US FDA has withdrawn approval due to a lack of efficacy of 17-OHPC for preterm birth (88), translational studies demonstrated that progesterone and 17-OHPC, specifically, may contribute to maternal immunomodulation and prevention of adverse perinatal outcomes other than preterm birth (41, 79). Furthermore, while nonselective COX inhibitors [i.e. non-steroidal anti-inflammatory drugs (NSAIDs)] are generally not recommend for use during pregnancy (89, 90), we have identified COX-1-dependent P4 regulation as a targetable pathway that warrants further investigation. Our results add to growing literature utilizing P4 to treat adverse pregnancy and perinatal outcomes (41, 79, 91, 92). With pregnant people considered an ‘at risk’ population for many infectious diseases ranging from influenza viruses and SARS-CoV-2 to Zika viruses, greater consideration should be given to identifying the mechanisms of adverse pregnancy outcomes and the off label use of compounds (e.g., 17-OHPC) that are safe during pregnancy and could mitigate the impact of infectious diseases at the maternal-fetal interface.

## 5 Acknowledgements

The authors would like to thank the Klein, Pekosz, Davis, and Baumgarth laboratories for discussions about these data. We would also like to thank the expert animal care staff at the Johns Hopkins School of Public Health for assistance with maintenance of pregnant dams.

## 7 Contributions

SK, PC, MP, and IB conceptualized and designed the experiments. PC, MP, JP, AC, and MS performed animal experiments. PC grew and quantified viruses and multiplexed cytokines. PC and MP performed ELISA and Western blot. PC, JP, and JinL performed flow cytometry. JunL stained, imaged, and analyzed placental tissue. PC statistically analyzed and graphed data. PC and SK wrote the manuscript with input from all authors. All authors read and provided edits to drafts and approved the final submission.

