## Supplemental Figures for "Suppression of progesterone by influenza A virus mediates adverse maternal and fetal outcomes in mice"

### Figure S1

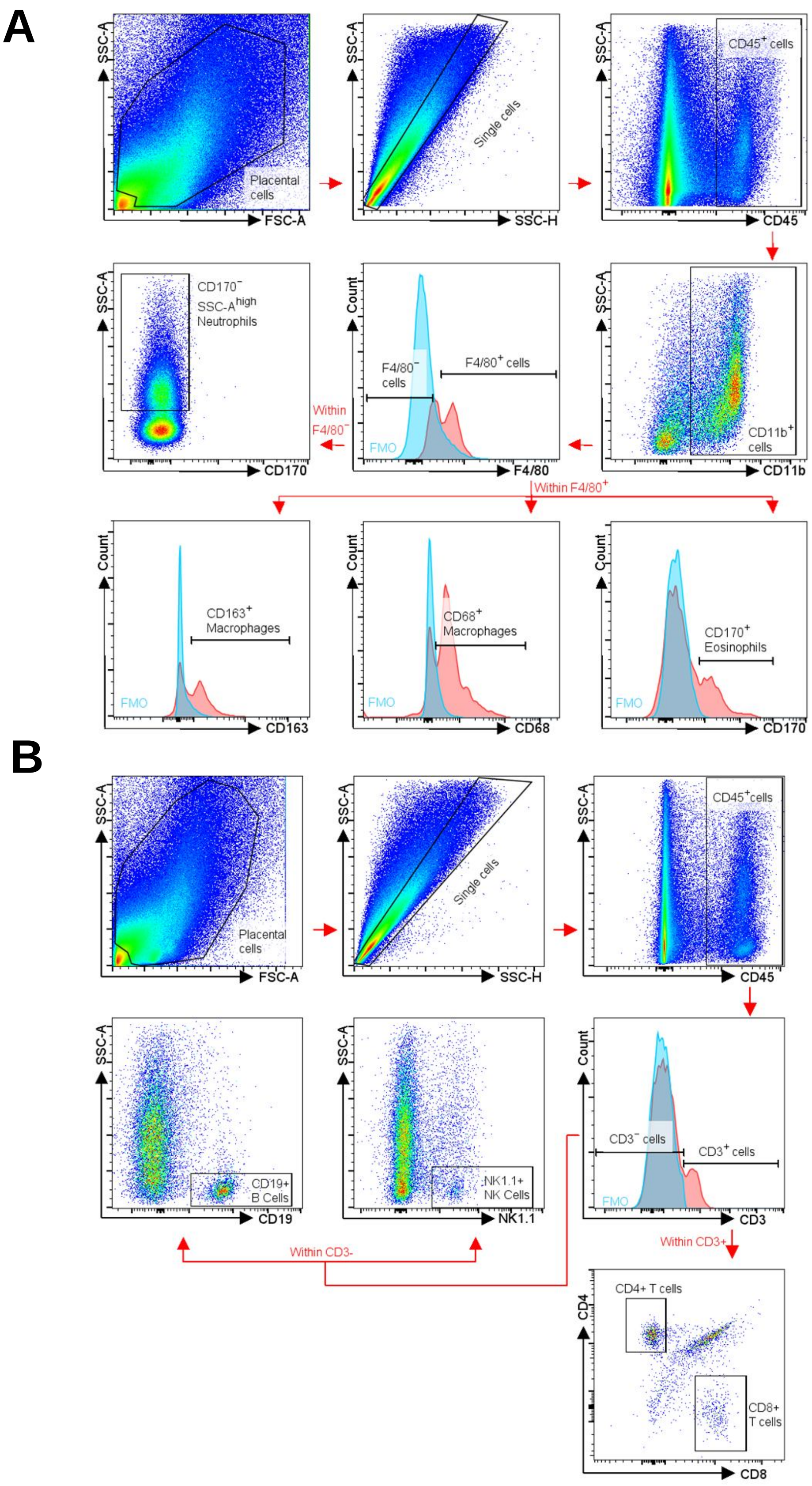


**Supplemental Figure 1. Gating strategy for immune cell phenotyping in placental tissue.**
Gating strategy for multi-color flow cytometry analysis of granulocyte and monocyte (A) as well as lymphocyte (B) immune cell populations. Debris and doublets were excluded by gating on side scatter height versus side scatter area to delaminate singlets, and specific populations including leukocytes, (CD45+), NK cells (CD45+, CD3-,CD19+), B cells (CD45+, CD3-,CD19+), CD4+ T cells (CD45+, CD3+, CD4+), CD8+ T cells (CD45+, CD3+, CD8a+), neutrophils (CD45+, CD11b+, F4/80-, CD170-, side scatter area high), eosinophils (CD45+, CD11b+, F4/80+, CD170+), CD163+ macrophages (CD45+, CD11b+, F4/80+, CD163+), and CD68+ macrophages (CD45+, CD11b+, F4/80+, CD68+) were gated based on unstained and fluorescent minus one (FMO) controls.

### Figure S2

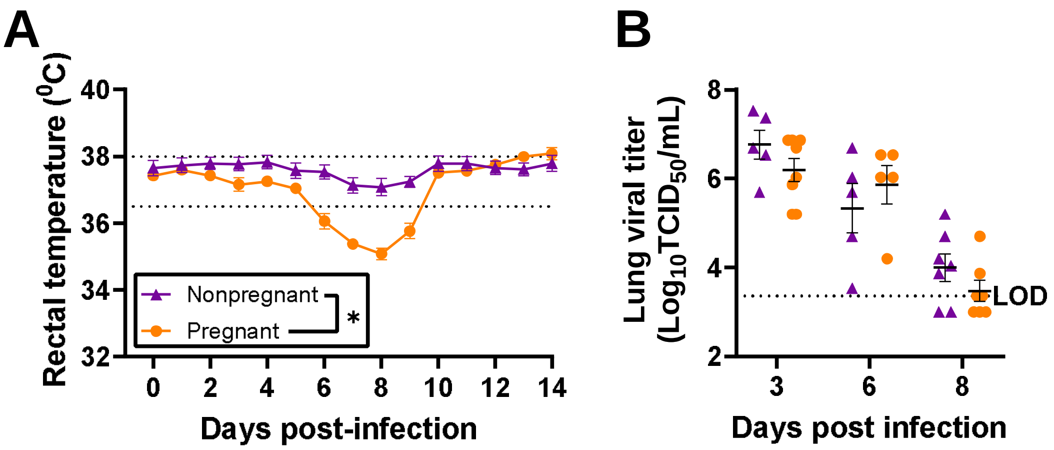


**Supplemental Figure 2. Pregnant mice experience more severe morbidity than nonpregnant females that is not mediated by lung viral load.**
Nonpregnant female mice or dams at embryonic day (E) 10 were intranasally inoculated with $10^3$ TCID50 2009 H1N1 or were mock inoculated with media. (A) Daily rectal temperatures were recorded as a measurement of morbidity through 14dpi. (B) Subsets of dams were euthanized to measure viral titers in lung tissue via TCID50 assay at 3 (E13), 6 (E16), and 8 (E18) days post infection. Individual symbols (A) or bars (B) represent the mean ± standard error of the mean per group from two independent replications (n =9-13/group for morbidity and 5-9/group for viral titers) with individual mice indicated by symbols (B). Significant differences ($p < 0.05$) were determined by two tailed unpaired t-test of AUCs (A) or two-way ANOVA with Bonferroni post hoc test (C) and are indicated by an asterisk (*).

### Figure S3

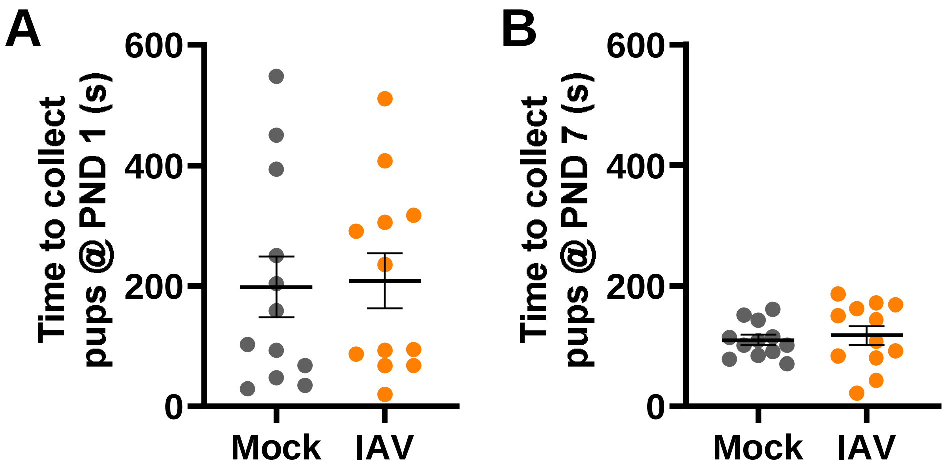


**Supplemental Figure 3. Influenza A virus infection during pregnancy does not impair postnatal pup retrieval.**

Dams at embryonic day (E) 10 were intranasally inoculated with $10^3$ TCID50 of 2009 H1N1 or were mock inoculated with media and were followed through delivery and to postnatal day (PND) 7. At PND1 (A) and PND7 (B), dams were subjected to a pup retrieval assay to assess postnatal care. Dams were placed in a clean cage for 10 min to acclimate, with pups removed and placed on a heating pad. 5 pups per dam were randomly distributed in the home cage away from the nest, and then the dams was returned to the home cage. The time in s it took for the dam to retrieve all 5 pups was recorded. Bars represent the mean ± standard error of the mean per group from two independent replications (n =12 dams/group) with or individual mice indicated by symbols. Significant differences ($p < 0.05$) were determined by two tailed unpaired t-test (A-D) and are indicated by an asterisk (*).

### Figure S4

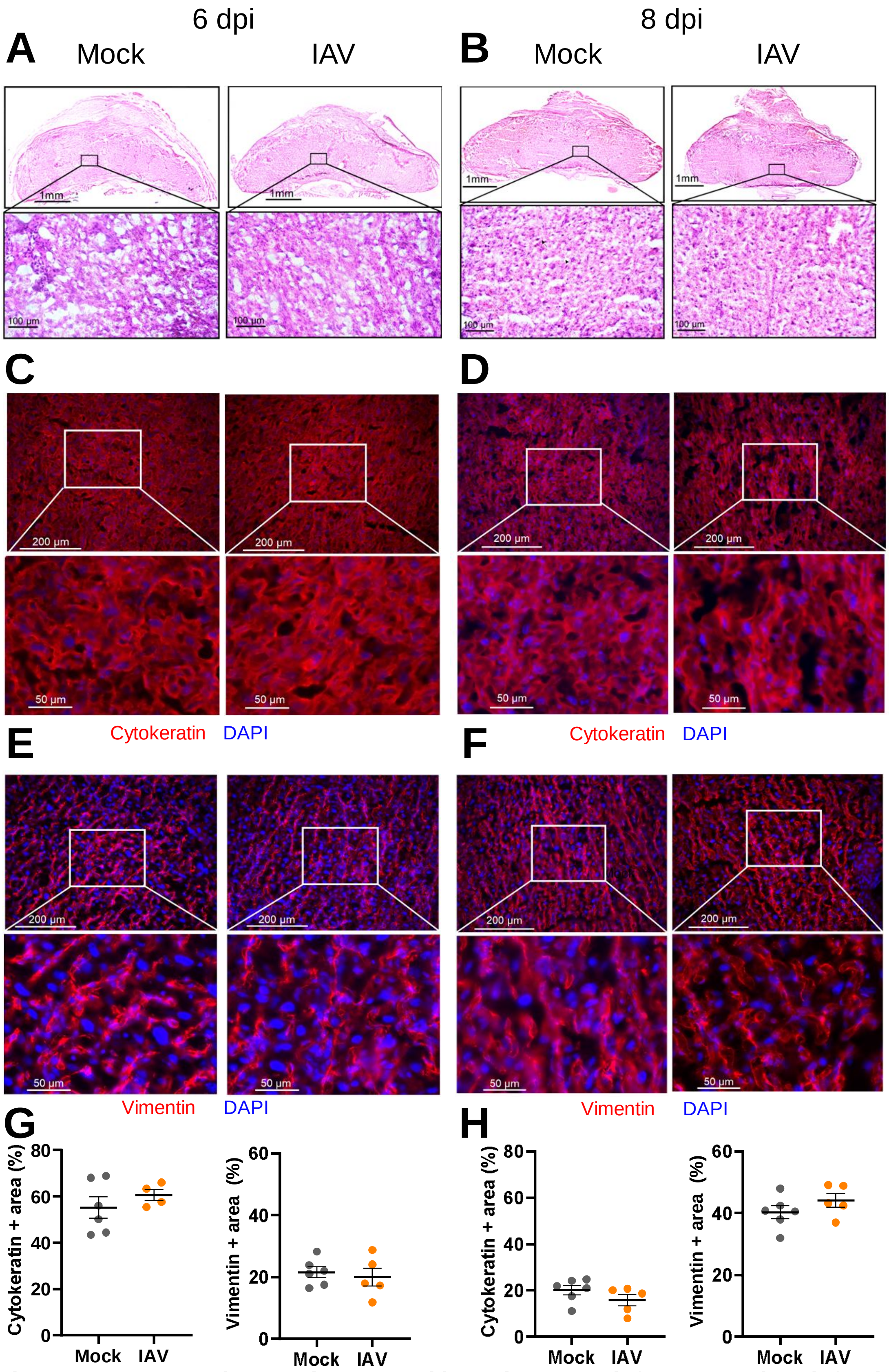


**Supplemental Figure 4. Placental damage is resolved by 6 days post influenza A virus infection.**
Dams at embryonic day (E) 10 were intranasally inoculated with $10^3$ $TCID_{50}$ of 2009 H1N1 or were mock inoculated with media, with placentas collected at 6 (A, C, E, G) and 8 (B, D, F, H) dpi. Representative H&E images were taken at 5x magnification and 20x magnification (A, B). Placentas were immunostained for cytokeratin (C, D, red) to mark trophoblasts or vimentin (E,F, red) to mark endothelial cells and DAPI (blue) to label nuclei. Representative images were taken at 20x magnification and further zoomed 2.9-fold. Quantification of the percentage positive area for each marker is shown (G-H). Bars represent the mean ± standard error of the mean (4-6/group) with each symbol indicating 1 placenta and is the mean quantification of 6 fields of view. Significant differences ($p < 0.05$) were determined by Mann-Whitney nonparametric test and are indicated by an asterisk (*). Scale bar: 1mm (A-B, upper panel/group), 100 µm (A-B, lower panel/group), 200 µm (C-F, upper panel/group), or 50 µm (C-F, lower panel/group)

### Figure S5

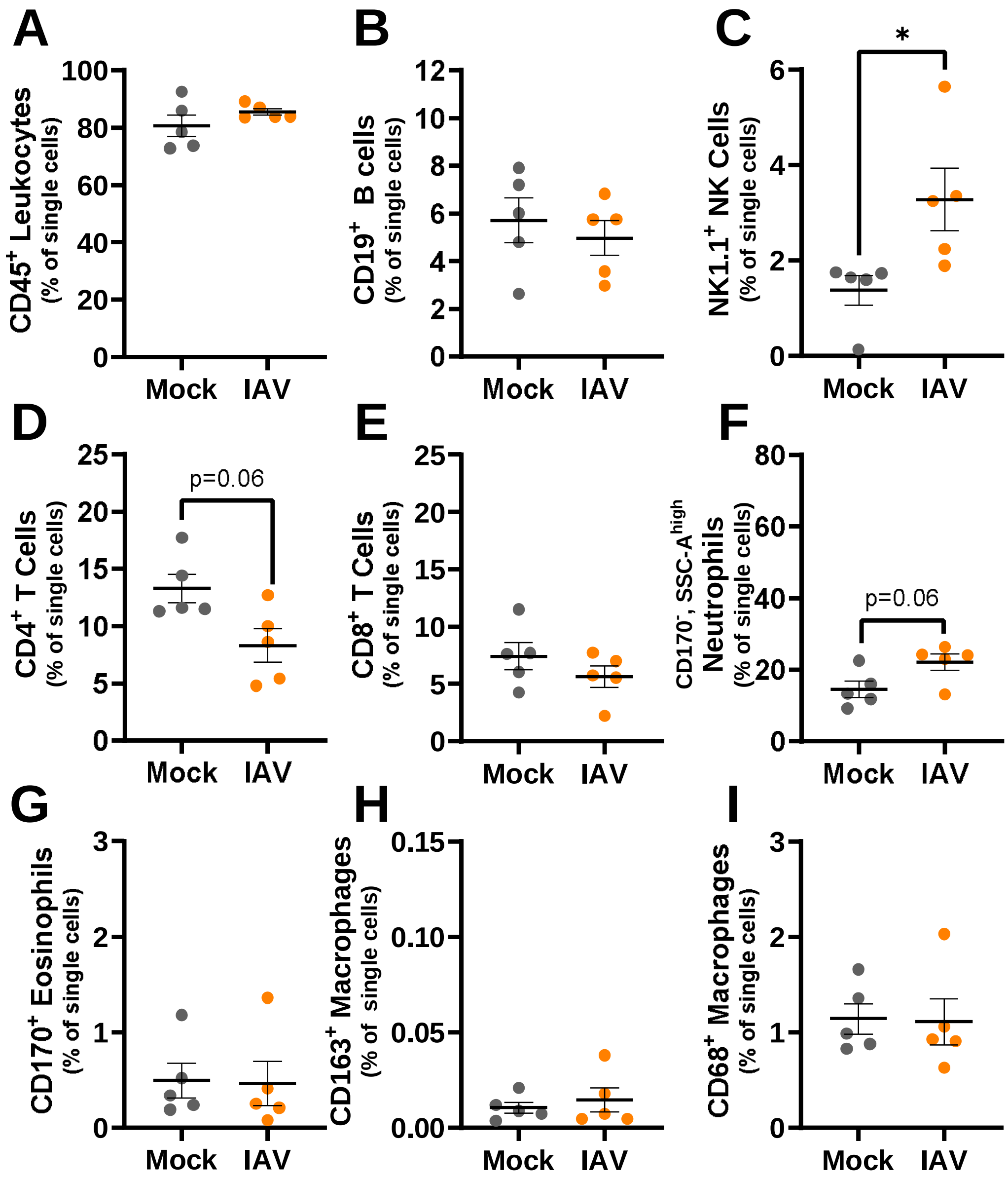


**Supplemental Figure 5. Immune cell frequencies in peripheral blood of dams infected with influenza A virus or mock inoculated.**
Dams at E10 were intranasally inoculated with $10^3$ TCID50 of 2009 H1N1 or were mock inoculated with media and were euthanized at 6 (E16) days post infection for flow cytometry to assess immune cell type frequencies in peripheral blood cells including leukocytes ($CD45^+$, A), NK cells ($CD45^+$, $CD3^-$,$CD19^+$, B), B cells ($CD45^+$, $CD3^-$,$CD19^+$, C), $CD4^+$ T-cells ($CD45^+$, $CD3^+$, $CD4^+$, D), $CD8^+$ T cells ($CD45^+$, $CD3^+$, $CD8a^+$, E), neutrophils ($CD45^+$, $CD11b^+$, $F4/80^-$, $CD170^-$, side scatter area high, F), eosinophils ($CD45^+$, $CD11b^+$, $F4/80^+$, $CD170^+$, G), $CD163^+$ macrophages ($CD45^+$, $CD11b^+$, $F4/80^+$, $CD163^+$, H), and CD68+ macrophages ($CD45^+$, $CD11b^+$, $F4/80^+$, $CD68^+$, I). Bars represent the mean ± standard error of the mean per group from two independent replications (n =5/group) with individual mice indicated by symbols. Significant differences ($p < 0.05$) were determined by Mann-Whitney nonparametric test and are indicated by an asterisk (*).

### Figure S6

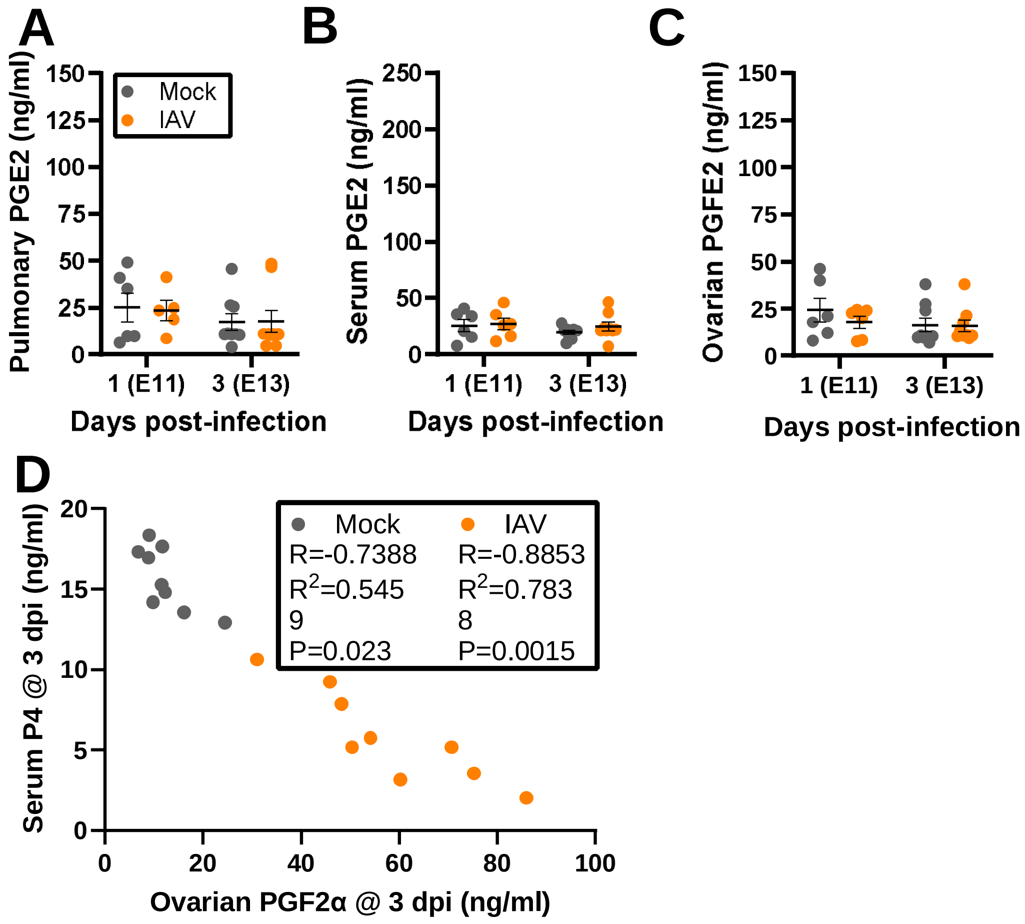


**Supplemental Figure 6. Influenza A virus infection does not alter concentrations of PGE2.**
Dams at embryonic day (E) 10 were intranasally inoculated with $10^3$ TCID50 of 2009 H1N1 or were mock inoculated with media. Dams were euthanized for tissue and serum collection at 1 (E11) or 3 (E13) days post infection. Concentrations of PGE2 (ng/ml) were measured by ELISA (A-C). Associations between serum P4 and ovarian PGF2α in dams IAV-infected or mock inoculated were analyzed by Pearson correlation analyses, with a significant association represented with the R statistic and associated P value (D). Data represent mean +/- standard error of the mean (n= 6-9/group). Significant differences (p<0.05) were determined by Kruskal–Wallis with Dunn's post hoc test

### Figure S7

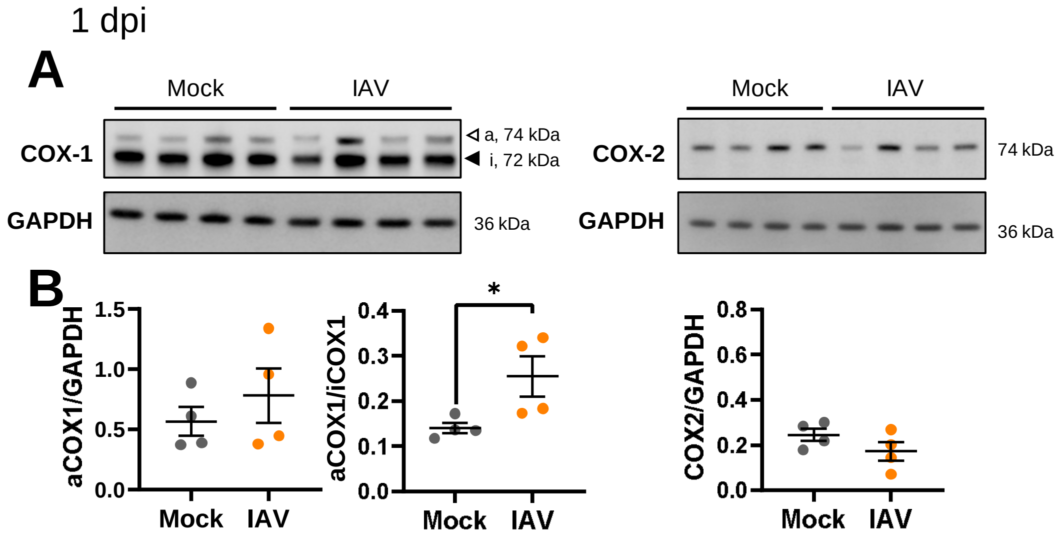


**Supplemental Figure 7. Active COX-1 as a ratio to inactive COX-1 is increased in the lungs at 1 dpi.**

Dams at E10 were intranasally inoculated with $10^3$ TCID5 of maH1N1 or were mock inoculated with media. COX-1 and COX-2 protein expression in lung tissue was assessed by western blot at 1dpi (A) and quantified as the fluorescence signal for each protein normalized to GAPDH. Active COX-1 is also shown as a ratio to inactive COX-1 (B). Lungs from two independent experiments were analyzed together to minimize interassay variability. Bars represent the mean ± standard error of the mean (n =4/group) with individual mice indicated by symbols. Significant differences ($p < 0.05$) were determined Mann-Whitney nonparametric test and are indicated by an asterisk (*).

### Figure S8

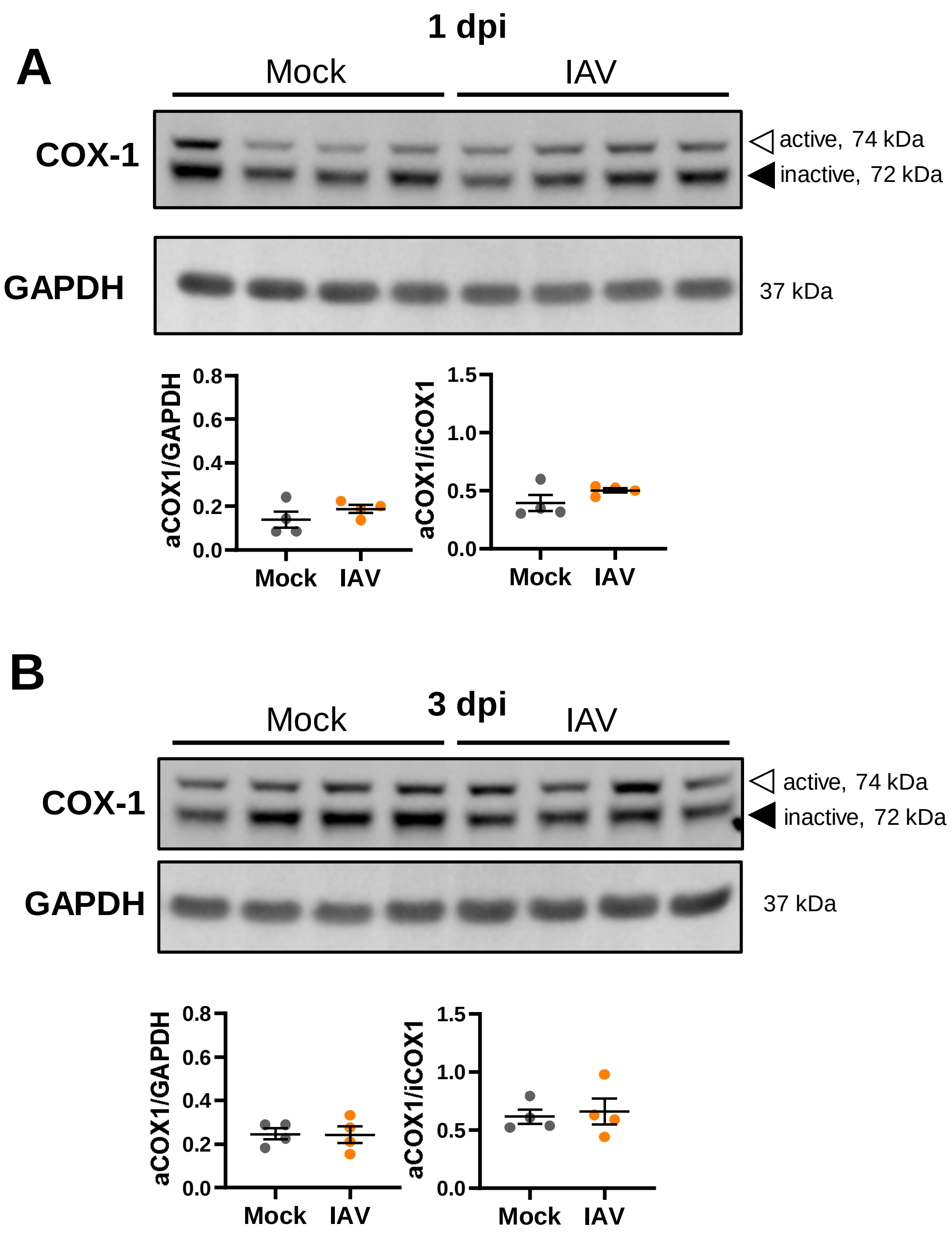


**Supplemental Figure 8. COX-1 expression is unchanged in the ovaries following maternal IAV infection**

Dams at E10 were intranasally inoculated with $10^3$ TCID50 of 2009 H1N1 or were mock inoculated with media. COX-1 and COX-2 protein expression in ovarian tissue was assessed by western blot at 1 (A) and 3 (B) dpi and quantified as the fluorescence signal for each protein normalized to GAPDH. Active COX-1 is also shown as a ratio to inactive COX-1 (B). Ovaries from two independent experiments were analyzed together to minimize interassay variability. Bars represent the mean ± standard error of the mean (n =4/group) with individual mice indicated by symbols. Significant differences ($p < 0.05$) were determined Mann-Whitney nonparametric test and are indicated by an asterisk (*).

### Figure S9

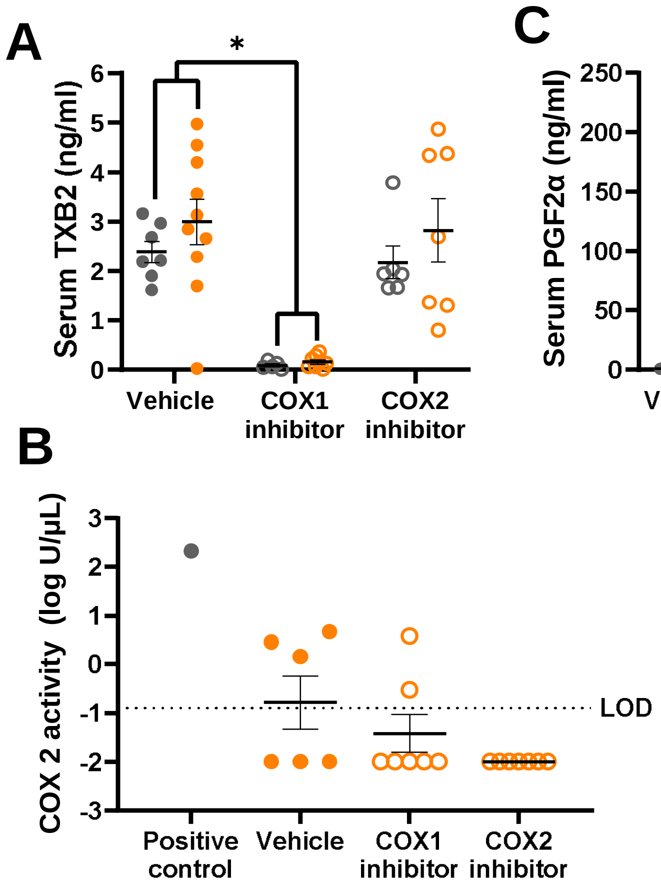


**Supplemental Figure 9. COX-1, but not COX-2 inhibition reduced serum PGF2α following influenza A virus infection**

Dams at embryonic day (E) 10 were intranasally inoculated with $10^3$ TCID50 of 2009 H1N1 or were mock inoculated with media. Dams were treated twice daily beginning 1 day before infection with either COX-1 or COX-2 inhibitor or vehicle until dams were euthanized at 3 days post infection, with for serum or lungs collected. Specificity of COX-1 inhibition was measured by inhibition of serum thromboxane B2 (ng/mL) by ELISA (A). Specificity of COX-2 inhibition was measured by COX-2 enzymatic activity in lung homogenate of IAV-infected dams. Serum concentrations PGF2α (ng/mL) was measured by ELISA (B). Significant differences ($p<0.05$) were determined by Kruskal–Wallis with Dunn's post hoc test and are indicated by an asterisk (*).

### Figure S10

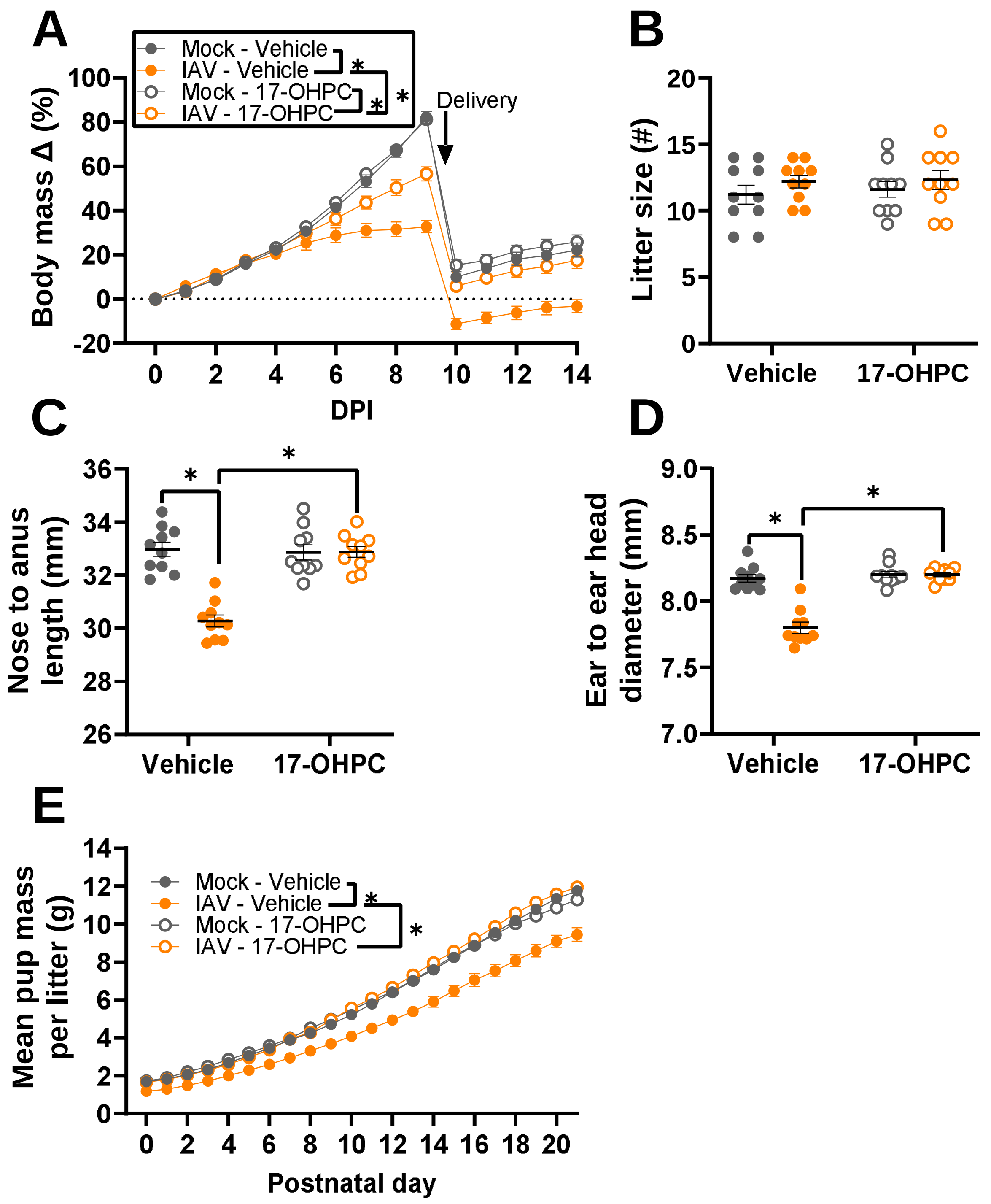


**Supplemental Figure 10. Synthetic progestin treatment improves maternal morbidity and prevents offspring growth restriction**

Dams at embryonic day (E) 10 were intranasally inoculated with $10^3$ TCID50 of 2009 H1N1 or were mock inoculated with media. At the time of inoculation, dams were subcutaneously administered 17α-hydroxyprogesterone caproate (17-OHPC, 2mg/dam). Body mass was measured daily as a measurement of maternal morbidity through 14dpi (A). Offspring were followed through postnatal day (PND) 21 to assess fetal outcomes. Litter size (B), nose to anus length (C), and ear to ear head diameter (D) were measured at PND 0. Average measurements of each independent litter were graphed to account for litter effects. Average pup mass for each litter was taken daily to track growth through PND 21 (E). Individual symbols (A, E) or bars (B-D) represent the mean ± standard error of the mean per group from two independent replications (n =10 mice or litters /group) with individual mice (A) or individual litter means (B-D) indicated by symbols. Significant differences ($p < 0.05$) were determined by two-way ANOVA with Bonferroni post hoc test of AUCs (A) or two-way ANOVA with Bonferroni post hoc test (B-E) and are indicated by an asterisk (*).

### Figure S11

## A

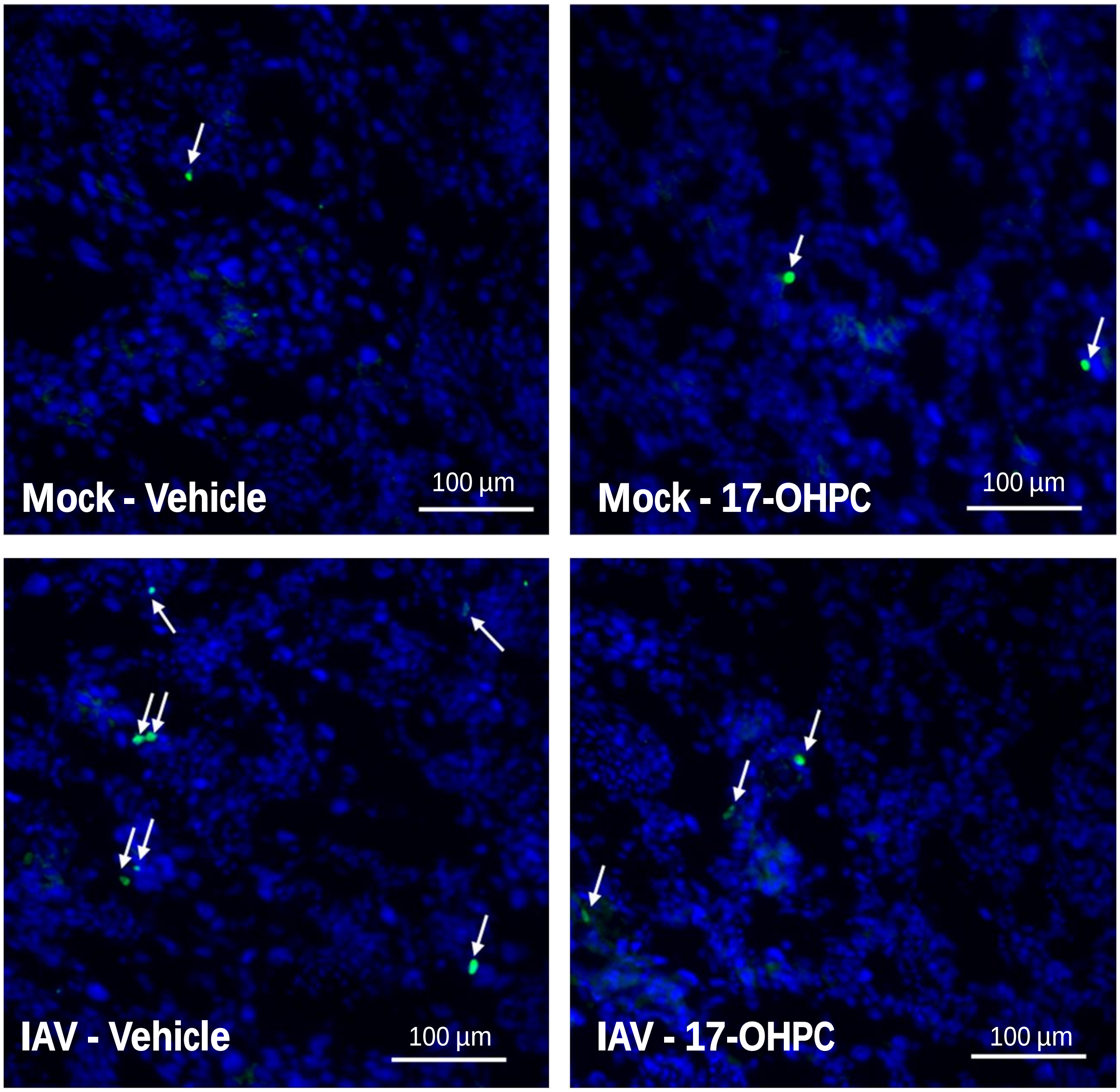


**Supplemental Figure 11. Synthetic progestin treatment reduces cell death in the placenta**
Dams at embryonic day (E) 10 were intranasally inoculated with $10^3$ TCID50 of 2009 H1N1 or were mock inoculated with media. At the time of inoculation, dams were subcutaneously administered 17α-hydroxyprogesterone caproate (17-OHPC, 2mg/dam). ). Placentas were collected at 3 (E13) days post infection (dpi). Representative TUNEL+ (green) stained placentas with DAPI to label nuclei taken (blue) at 20x magnification with TUNEL+ nuclei indicated with arrows. Scale bar: 100 µm
