## Supplemental Text 1 for "Suppression of progesterone by influenza A virus mediates adverse maternal and fetal outcomes in mice"

**Text S1: Experimental Details for miniaturized multiplex cytokine analysis**

Briefly, 96-well DA-Bead Plates (Curiox 96-CC-BD-05) were blocked with 10 µL per well of 1% BSA (Sigma A9647) in PBS for 30 min in a humidified magnet box (Curiox HMBXWT01) at room temperature on a rotator. The plate was washed with Curiox wash buffer [0.1% BSA (Sigma A9647) and 0.05% Tween-20 (Sigma P1379)] on a DropArray LT210 (Curiox). Procartaplex beads (5 µL) were added to each well, followed by 5 µL of serially diluted ProCartaplex standards or 10 µL tissue homogenate per well, with each sample assayed in duplicate. The plate was incubated overnight in a humidified magnet box on a rotator at 4°C. The following day, the plate was washed three times, and 5 µL of ProCartaPlex detection antibody added per well. The plate was incubated for 1 h in a humidified magnet box on a rotator. After incubation, 10 µL streptavidin-phycoerythrin was added to each well and incubated for 30 min in a humidified magnet box on a rotator. The plate was washed three times, and the beads were reconstituted in Curiox wash buffer and transferred to a skirted v-bottom 96 well plate (Biorad HSP9601) for reading on a MagPix system (Luminex). Analysis was performed using BioPlex Manager (version 6.2). The following analytes were quantified: G-CSF, GM-CSF, M-CSF, CXCL1, CXCL2, CXCL5, CXCL10, CCL2, CCL3, CCL4, CCL5, CCL7, CCL11, IFNα, IFNγ, IL-1β, IL-1α, IL-2, IL-3, IL-2R, IL-4, IL-5, IL-6, IL-7, IL-7Rα, IL-9, IL-10, IL-12p70, IL-13, IL-15/IL-15R, IL-17A IL-18, IL-19, IL-22, IL-23, IL-25, IL-27, IL-28, IL-31, IL-33, IL-33R, Leukemia inhibitory factor (LIF), TNFα, receptor activator of nuclear factor kappa-B ligand (RANKL), B-cell activating factor (BAFF), Betacellulin, vascular endothelial growth factor (VEGF-A), and Leptin.
