## Supplemental Table 1 for "Suppression of progesterone by influenza A virus mediates adverse maternal and fetal outcomes in mice"

Table S1.

| Antibodies used for Flow Cytometry, Western Blot, and Immunohistochemistry | | | | | |
| --- | --- | --- | --- | --- | --- |
| Use | Antigen | Fluorophore | Clone | Catalog # | Dilution |
| Flow Cytometry Panel 1 | CD45 | Super Bright 600 | 30-F11 | 63-0451-82 | 1:100 |
|  | CD11b | PerCP-Cyanine5.5 | M1-70 | 45-0112-82 | 1:100 |
|  | F4/80 | Efluor 450 | BM8 | 48-4801-82 | 1:100 |
|  | CD170 | PE-Cyanine7 | 1RNM44N | 25-1702-82 | 1:100 |
|  | CD163 | PE | TNKUPJ | 12-1631-82 | 1:100 |
|  | CD168 | Alexa Fluor 488 | FA-11 | 53-0681-82 | 1:100 |
| Flow Cytometry Panel 2 | CD45 | PE | 30-F11 | 12-0451-82 | 1:100 |
|  | NK1.1 | PE-Cyanine7 | PK136 | 25-5941-82 | 1:100 |
|  | CD3 | eFluor 450 | 17A2 | 48-0032-82 | 1:100 |
|  | CD8a | eFluor 506 | 53-6.7 | 69-0081-82 | 1:100 |
|  | CD19 | Super Bright 600 | eBio1D3 | 63-0193-82 | 1:100 |
| Western Blot 1° | COX-1 | Unconjugated | EPR5866 | ab109025 | 1:1000 |
|  | COX-2 | Unconjugated | EPR12012 | ab179800 | 1:1000 |
|  | GAPDH | Unconjugated | 6C5 | ab8245 | 1:10,000 |
| Western Blot 2° | Mouse IgG | Alexa Fluor 488 | Polyclonal | A11001 | 1:10,000 |
|  | Rabbit IgG | Alexa Fluor 647 | Polyclonal | A32795 | 1:10,000 |
| Immunohistochemistry 1° | Vimentin | Unconjugated | EPR3776 | ab92547 | 1:200 |
|  | Cytokeratin | Unconjugated | Polyclonal | Z0622 | 1:200 |
| Immunohistochemistry 2° | Rabbit IgG | Alexa Fluor 594 | Polyclonal | R37119 | 1:500 |
