## Supplemental Table 2 for "Suppression of progesterone by influenza A virus mediates adverse maternal and fetal outcomes in mice"

|  | Concentration in lung (pg/ml) | | | | | |
| --- | --- | --- | --- | --- | --- | --- |
|  | 3 DPI | | 6 DPI | | 8 DPI | |
| Cytokine | Mock | IAV | Mock | IAV | Mock | IAV |
| IFNα | **313.76 ± 22.78** | **1592.39 ± 238.61** | **32.75 ± 2.24** | **3073.36 ± 780.68** | 269.54 ± 40.96 | 264.36 ± 33.29 |
| IFNγ | **5.53 ± 0.66** | **38.38 ± 3.18** | **42.21 ± 11.32** | **2825.62 ± 555.62** | **3.46 ± 0.65** | **95.81 ± 14.03** |
| IL-28 | **6249.17 ± 844.49** | **48262.9 ± 4481.6** | **5322.15 ± 1441.71** | **31463.9 ± 3956.8** | **4038.66 ± 882.01** | **22291.2 ± 3079.6** |
| IL-1α | **150.09 ± 15.54** | **1487.63 ± 43.51** | 391.03 ± 120.85 | 337.19 ± 48.59 | 134.76 ± 26.61 | 142.7 ± 23.11 |
| IL-1β | **49.55 ± 5.54** | **85.98 ± 2.43** | 22.17 ± 4.14 | 21.33 ± 4.84 | 27.67 ± 6.4 | 29.66 ± 6.24 |
| IL-18 | **144.67 ± 13.59** | **2292.77 ± 297.97** | 438.83 ± 129.84 | 1027.87 ± 126.7 | 173.54 ± 33.54 | 272.93 ± 21.9 |
| IL-33 | **21315.8 ± 2033.2** | **42072.2 ± 1753.8** | 17261.6 ± 3577.7 | 18741.6 ± 3967.1 | 22520.9 ± 2530 | 26217.6 ± 3201.5 |
| IL-33R | 1797.38 ± 161.98 | 2251.6 ± 242.43 | 165.43 ± 45.11 | 221.19 ± 52.13 | 1845.34 ± 232.88 | 1935.45 ± 207.56 |
| IL-2 | **145.43 ± 3.63** | **82.15 ± 6.01** | 774.93 ± 57.72 | 574.3 ± 136.31 | 152.14 ± 5.31 | 132.36 ± 11.41 |
| IL-2R | **6.26 ± 0.38** | **7.93 ± 0.31** | 38.4 ± 7.21 | 44.88 ± 5.11 | **6.08 ± 0.61** | **14.11 ± 1.04** |
| IL-4 | **61.92 ± 4.82** | **164.43 ± 17.86** | 200.15 ± 37.85 | 321.39 ± 89.33 | 80.14 ± 5.91 | 78.21 ± 5.7 |
| IL-7 | 108.23 ± 21.47 | 171.83 ± 4.03 | 221.54 ± 41.24 | 233.16 ± 83.24 | 177.04 ± 17.71 | 144.45 ± 21.28 |
| IL-7Rα | 205.21 ± 6.14 | 213.96 ± 10.63 | 2036.02 ± 278.65 | 1511.57 ± 267.38 | 232.94 ± 14.6 | 289.63 ± 31.77 |
| IL-9 | 74.97 ± 0.86 | 309.42 ± 44.27 | 109.35 ± 20.4 | 114.23 ± 5.1 | 75.15 ± 1.53 | 76.57 ± 1.98 |
| IL-15/ IL-15R | 3.25 ± 0.25 | 3.33 ± 0.16 | 63.73 ± 21.3 | 52.03 ± 10.11 | 3.32 ± 0.28 | 3.56 ± 0.23 |
| TNFα | **7.3 ± 0.62** | **188.6 ± 20.79** | **49.64 ± 7.89** | **180.2 ± 14** | **6.41 ± 0.65** | **62.84 ± 7.95** |
| RANKL | 20.6 ± 1.33 | 45.08 ± 6.87 | 20.44 ± 3.66 | 13.03 ± 3.88 | 21.69 ± 1.9 | 34.92 ± 8.57 |
| BAFF | 81.73 ± 9.34 | 89.92 ± 9.15 | **49.68 ± 13.66** | **394.01 ± 84.01** | **62.91 ± 12.74** | **205.94 ± 21.71** |
| IL-10 | **87.4 ± 14.52** | **272.04 ± 9.53** | **1819.56 ± 156.46** | **4879.26 ± 770.32** | 57.11 ± 18.85 | 32.18 ± 6.26 |
| IL-19 | **2160.45 ± 193.39** | **3409.97 ± 144.92** | 1298.46 ± 267.43 | 8410.21 ± 2057.18 | 1517.26 ± 256.38 | 1465.48 ± 145.73 |
| IL-22 | **180.59 ± 10.72** | **141.8 ± 3.7** | **134.5 ± 14.78** | **624.01 ± 53.89** | 158.15 ± 16.86 | 135.55 ± 11.51 |
| IL-12p70 | **6.87 ± 0.86** | **13.09 ± 1.44** | 154.73 ± 61.72 | 97.41 ± 49.24 | 7.83 ± 1.41 | 6.65 ± 1.04 |
| IL-23 | 1917.25 ± 107.75 | 1478.44 ± 96.92 | 504.56 ± 45.97 | 722.49 ± 281.48 | 1536.83 ± 267.31 | 1343.24 ± 238.35 |
| IL-27 | **24.73 ± 1.75** | **236.49 ± 28.93** | **63.7 ± 9.01** | **442 ± 48.58** | **20.19 ± 2.09** | **225.28 ± 48.77** |
| IL-17A | 258.27 ± 21.66 | 212.48 ± 17.76 | 441.07 ± 105.38 | 567.71 ± 91.25 | 158.32 ± 22.41 | 133.75 ± 11.69 |
| IL-25 | 138.84 ± 22.49 | 118.4 ± 18.79 | 199.81 ± 52.74 | 493.25 ± 154.37 | 116.14 ± 29.34 | 168.51 ± 39.49 |
| IL-6 | **459.16 ± 69.11** | **2587.66 ± 270.99** | **159.39 ± 39.07** | **1067.93 ± 193.85** | 721.35 ± 164.48 | 1075.46 ± 276.02 |
| IL-31 | **1196.83 ± 40.12** | **1453.67 ± 61.29** | **2545.94 ± 67.57** | **2165.08 ± 71.92** | 1464.61 ± 100.64 | 1406.68 ± 96.51 |
| LIF | **71.99 ± 3.6** | **435.7 ± 23.3** | 163.93 ± 32.26 | 202.6 ± 39.76 | **79.53 ± 7.05** | **232.25 ± 21.1** |
| CCL2 | **355.68 ± 50.17** | **4228.42 ± 497.72** | **50.16 ± 9.49** | **1915.85 ± 341.04** | **190.33 ± 48.34** | **4205.22 ± 669.41** |
| CCL3 | **4.85 ± 0.64** | **135.47 ± 19.61** | **4.06 ± 0.9** | **58 ± 9.14** | **6.02 ± 1.58** | **236.63 ± 36.47** |
| CCL4 | **13.05 ± 1.59** | **201.69 ± 18.89** | **16.18 ± 2.27** | **112.31 ± 17.49** | **10.16 ± 1.29** | **292.47 ± 40.86** |
| CCL5 | **64.29 ± 7.09** | **214.56 ± 34.96** | 139.72 ± 10.8 | 119.78 ± 40.35 | **76.13 ± 14.57** | **219.91 ± 13.34** |
| CCL7 | **92.05 ± 10.02** | **1537.52 ± 219** | **118.08 ± 23.26** | **513.9 ± 46.97** | **83.37 ± 15.65** | **428.93 ± 52.97** |
| CCL11 | **257.39 ± 15.34** | **3470.88 ± 242.97** | 1101.51 ± 204.96 | 788.67 ± 178.23 | 260.78 ± 17.64 | 869.8 ± 157.55 |
| CXCL1 | **26.09 ± 2.59** | **1256.67 ± 129.62** | 107.47 ± 17.08 | 70.96 ± 17.62 | **21.95 ± 4.7** | **181.88 ± 29.25** |
| CXCL2 | **9.01 ± 0.4** | **148.04 ± 11.13** | 195.41 ± 62.09 | 247.81 ± 31.45 | 9.82 ± 0.88 | 9.77 ± 0.82 |
| CXCL5 | **389.45 ± 39.56** | **1635.92 ± 235.33** | 1095.4 ± 350.61 | 1117.67 ± 301.45 | 470.01 ± 63.01 | 488.17 ± 71.68 |
| CXCL10 | 148.68 ± 12.14 | 175.13 ± 13.31 | 223.16 ± 40.11 | 656.16 ± 134.24 | **130.67 ± 20.12** | **1202.48 ± 140.56** |
| BTC | 42.3 ± 4.36 | 40.88 ± 5.5 | 241.25 ± 51.93 | 290.41 ± 74.93 | 29.62 ± 7.57 | 30.91 ± 6.56 |
| G-CSF | 24.89 ± 1.3 | 25.24 ± 2.53 | 206.25 ± 56.08 | 279.64 ± 48.42 | 21.17 ± 2.19 | 21.72 ± 2.02 |
| GM-CSF | 34.49 ± 1.41 | 31.49 ± 1.73 | **219.3 ± 88.03** | **1384.27 ± 221.63** | 39.05 ± 2.89 | 38.36 ± 2.77 |
| M-CSF | 7.61 ± 0.99 | 9.61 ± 1.43 | 30.04 ± 5.05 | 11.61 ± 2.62 | 1.73 ± 1.11 | 1.01 ± 0.4 |
| IL-3 | 5.57 ± 0.36 | 5.97 ± 0.58 | 43.42 ± 8.9 | 49.62 ± 8.93 | 2.63 ± 0.74 | 3.99 ± 1.38 |
| VEGF-A | 759.17 ± 21.46 | 1149.2 ± 223.96 | 3482.35 ± 504.77 | 2619.89 ± 438.37 | 637.16 ± 29.91 | 741.89 ± 48.49 |
| IL-13 | 15.55 ± 1.9 | 17.27 ± 1.72 | 68.54 ± 12.33 | 64.61 ± 25.2 | 11.44 ± 2.1 | 13.5 ± 2.26 |
| IL-5 | **51.22 ± 5.43** | **208.27 ± 13.6** | 435.97 ± 75.53 | 237.14 ± 54.61 | 79.7 ± 14.91 | 109.76 ± 12.49 |
| Leptin | 147 ± 14.47 | 259.2 ± 42.94 | 146.98 ± 35.63 | 210.54 ± 32.98 | 140.35 ± 30.66 | 133.35 ± 30.04 |

Table S2.

* Bold indicates a statistically significant difference between mock and IAV at a given DPI based on multiple unpaired t tests with Holm-Šídák correction for multiple comparisons, adjusted p value <0.05 (n=12-14/group)
