## Supplemental Table 3 for "Suppression of progesterone by influenza A virus mediates adverse maternal and fetal outcomes in mice"

|  | Concentration in spleen (pg/ml) | | | | | |
| --- | --- | --- | --- | --- | --- | --- |
|  | 3 DPI | | 6 DPI | | 8 DPI | |
| Cytokine | Mock | IAV | Mock | IAV | Mock | IAV |
| IFNα | 475.13 ± 65.44 | 535.7 ± 62.38 | 2236.24 ± 572.55 | 2648.55 ± 681.74 | 415.65 ± 14.34 | 425.06 ± 24.92 |
| IFNγ | 11.09 ± 1.88 | 12.58 ± 1.29 | 256.05 ± 60.83 | 515.28 ± 100.72 | **8.76 ± 1.52** | **225.23 ± 49.61** |
| IL-28 | 3666.45 ± 822.28 | 5988.05 ± 1725.7 | 32405.03 ± 4339.7 | 34705.8 ± 3363.33 | 8833.65 ± 2194.48 | 11144.7 ± 2398.02 |
| IL-1α | 394.39 ± 91.81 | 340.79 ± 71.9 | 521.35 ± 108.37 | 636.05 ± 56.81 | 140.09 ± 28.11 | 170.05 ± 25.39 |
| IL-1β | 36.71 ± 5.86 | 41.94 ± 6.12 | 203.08 ± 34.89 | 191.79 ± 44.64 | 66.09 ± 10.84 | 78.37 ± 18.22 |
| IL-18 | 1245.51 ± 370.32 | 1184.5 ± 327.13 | 1957.12 ± 323.33 | 2343.25 ± 325.65 | 770.3 ± 84.77 | 923.56 ± 129.89 |
| IL-33 | 15483.36 ± 1927.9 | 22102 ± 1956.02 | 19867.3 ± 3931.06 | 17948.65 ± 3254.2 | 36534.19 ± 3217.7 | 40370.07 ± 2758.3 |
| IL-33R | 744.75 ± 167.69 | 520.02 ± 114.51 | 11503.9 ± 2280.9 | 10662.33 ± 2263.5 | 3756.87 ± 493.09 | 3126.47 ± 657.15 |
| IL-2 | 134.27 ± 15.05 | 177.61 ± 9.37 | 598.86 ± 90.58 | 609.21 ± 133.21 | 174.48 ± 18.25 | 173.28 ± 8.69 |
| IL-2R | 31.5 ± 6.07 | 29.81 ± 5.3 | 58.86 ± 7.3 | 61.06 ± 4.44 | 12.34 ± 1.51 | 14.09 ± 1.77 |
| IL-4 | 117.21 ± 19.61 | 135.79 ± 23.06 | 358.33 ± 68.65 | 311.4 ± 63.46 | 90.53 ± 1.42 | 90.83 ± 1.23 |
| IL-7 | 351.36 ± 46.59 | 360.91 ± 34.91 | 746.21 ± 186.83 | 774.71 ± 194.68 | 431.62 ± 71.26 | 397.85 ± 64.88 |
| IL-7Rα | 640.7 ± 192.62 | 718.32 ± 141.56 | 2444.38 ± 516.24 | 2599.42 ± 512.94 | 976.8 ± 158.04 | 915.5 ± 158.25 |
| IL-9 | 145.01 ± 26.43 | 83.02 ± 1.17 | 183.43 ± 18.34 | 161.13 ± 13.51 | 113.56 ± 28.34 | 158.19 ± 32.23 |
| IL-15/ IL-15R | 149.56 ± 31.9 | 135.81 ± 27.42 | 74.45 ± 2.86 | 269.68 ± 67.09 | 77.39 ± 26.19 | 85.84 ± 23.48 |
| TNFα | 56.07 ± 6.31 | 60.77 ± 6.82 | 220.34 ± 41.05 | 182.91 ± 28.45 | **83.9 ± 12.73** | **2622.37 ± 568.78** |
| RANKL | 43.62 ± 4.77 | 38.63 ± 2.96 | 24.66 ± 5.62 | 24.54 ± 6.37 | 102.85 ± 10.1 | 113.2 ± 6.88 |
| BAFF | 584.96 ± 65.4 | 438.38 ± 92.77 | 242.72 ± 49.71 | 286.79 ± 90.1 | 581.6 ± 81.76 | 554.27 ± 105.3 |
| IL-10 | 324.68 ± 48.68 | 370.01 ± 51.64 | 4681.29 ± 878.08 | 4100.14 ± 859.22 | 155.09 ± 45.12 | 149.16 ± 43.51 |
| IL-19 | 3928.39 ± 348.59 | 3344.45 ± 336.62 | 7232.48 ± 1778.37 | 8604.63 ± 2123.97 | 2060.88 ± 612.15 | 1786.72 ± 223.98 |
| IL-22 | 119.44 ± 12.22 | 162.59 ± 26.87 | 259.14 ± 45.3 | 320.95 ± 83.75 | 376.09 ± 98.5 | 478.64 ± 90.95 |
| IL-12p70 | 16.26 ± 2.24 | 16.88 ± 1.27 | 233.86 ± 55.57 | 135.32 ± 43.86 | 18.81 ± 3.64 | 22.83 ± 5.99 |
| IL-23 | 2848.88 ± 154.03 | 2304.25 ± 110.07 | 426.91 ± 59.36 | 734.46 ± 300.58 | 2329.83 ± 225.73 | 2827.06 ± 77.72 |
| IL-27 | 61.08 ± 8.67 | 69.98 ± 9.7 | 988.41 ± 175.4 | 792.64 ± 180.05 | 108.38 ± 7.93 | 149.01 ± 41.23 |
| IL-17A | 139.36 ± 53.65 | 112.04 ± 21.49 | 1101.98 ± 273.03 | 2261.81 ± 303.83 | 295.81 ± 37.57 | 306.08 ± 32.75 |
| IL-25 | 206.45 ± 51.8 | 189.84 ± 45.3 | 354.4 ± 76.44 | 410.4 ± 84.48 | 261.68 ± 51.53 | 330.73 ± 35.18 |
| IL-6 | 1663.75 ± 252.26 | 1943.53 ± 390.42 | 665.19 ± 176.88 | 760.33 ± 126.31 | 1578.02 ± 266.88 | 2257.98 ± 417.17 |
| IL-31 | 1443.47 ± 31.38 | 1450.11 ± 27.1 | 1735.55 ± 218.61 | 1682.15 ± 189.6 | 1726.63 ± 21.2383 | 1875.48 ± 82.05 |
| LIF | 54.24 ± 6.4 | 43.8 ± 6.65 | 180.37 ± 27.34 | 240.67 ± 26.69 | 103.13 ± 11.84 | 126.44 ± 18.36 |
| CCL2 | 590.16 ± 29.45 | 598.89 ± 25.77 | **909.51 ± 155.68** | **2420.58 ± 293.03** | 370.39 ± 92.88 | 513.99 ± 121.9 |
| CCL3 | 23.09 ± 4.75 | 33.57 ± 1.34 | 47.49 ± 5.4 | 36.7 ± 5.92 | 28.5 ± 3.02 | 43.92 ± 4.32 |
| CCL4 | 41.45 ± 10.4 | 42.06 ± 8.87 | **49.8 ± 7.33** | **89.47 ± 5.79** | 20.81 ± 5.03 | 44.63 ± 8.69 |
| CCL5 | 194.98 ± 6.06 | 155.64 ± 12.64 | 89.68 ± 16.17 | 200.26 ± 42.16 | 248.37 ± 47.73 | 170.87 ± 16.79 |
| CCL7 | 219.71 ± 24.22 | 283.1 ± 6.96 | **212.2 ± 46.93** | **513.71 ± 37.68** | 129.99 ± 21.76 | 186.29 ± 17.57 |
| CCL11 | 1231.61 ± 78.96 | 1320.29 ± 88.08 | 657.72 ± 74.66 | 512.62 ± 69.9 | 262.6 ± 72.33 | 344.5 ± 72.81 |
| CXCL1 | 120.03 ± 14.46 | 125.07 ± 9.57 | 1398.82 ± 331.04 | 1079.74 ± 357.92 | 74.11 ± 13 | 81.7 ± 10.77 |
| CXCL2 | 71.39 ± 1.57 | 64.14 ± 4.59 | 169.35 ± 52.7 | 126.77 ± 18.46 | 34.73 ± 7.88 | 25.05 ± 5.62 |
| CXCL5 | 1562.1 ± 120.99 | 1378.77 ± 95.85 | 3328.64 ± 93.48 | 3268.39 ± 79.77 | 1846.27 ± 295.18 | 2306.54 ± 339.17 |
| CXCL10 | 234.04 ± 17.01 | 214.37 ± 17.26 | 457.29 ± 75.55 | 536.61 ± 87.39 | 307.59 ± 30.71 | 356.58 ± 40.95 |
| BTC | 402.75 ± 38.75 | 375.48 ± 39.18 | 463.28 ± 103.22 | 533.78 ± 74.92 | 93.07 ± 18.07 | 75.49 ± 13.58 |
| G-CSF | 25.48 ± 4.82 | 26.38 ± 5.14 | 233.48 ± 49.12 | 404.13 ± 133 | 31.39 ± 2.596 | 49.1 ± 13.97 |
| GM-CSF | 241.48 ± 21.5 | 303.76 ± 30.73 | 1716.07 ± 347.77 | 1751.73 ± 282.34 | 162.33 ± 78.63 | 155.45 ± 67.71 |
| M-CSF | 15.38 ± 1.42 | 14.26 ± 1.08 | 43.14 ± 7.25 | 39.95 ± 6.8 | 31.06 ± 7.05 | 28.28 ± 5.04 |
| IL-3 | 7.5 ± 1.67 | 6.94 ± 1.47 | **28.85 ± 7.05** | **100.87 ± 10.69** | 20.34 ± 5.63 | 26.98 ± 4.64 |
| VEGF-A | 169.86 ± 51.95 | 224.9 ± 57.93 | 3481.27 ± 542.81 | 3000.36 ± 461.12 | 405.76 ± 67.75 | 427.69 ± 58.09 |
| IL-13 | 55.05 ± 11.73 | 66.16 ± 11.16 | 163.66 ± 30.98 | 141.7 ± 32.13 | 64.74 ± 13.15 | 73.81 ± 13.08 |
| IL-5 | 193.83 ± 30.11 | 231.08 ± 12.07 | 201.41 ± 43.26 | 260.56 ± 65.9 | 171.03 ± 35.24 | 127.95 ± 23.71 |
| Leptin | 833.34 ± 157.71 | 883.13 ± 159.05 | 249.85 ± 100.03 | 285.45 ± 96.86 | 2712.18 ± 1104.7 | 1849.78 ± 763.55 |

Table S3.

Bold indicates a statistically significant difference between mock and IAV at a given DPI based on multiple unpaired t tests with Holm-Šídák correction for multiple comparisons, adjusted p value <0.05 (n=12-14)
