## Supplemental Table 4 for "Suppression of progesterone by influenza A virus mediates adverse maternal and fetal outcomes in mice"

|  | Concentration in placenta (pg/ml) | | | | | |
| --- | --- | --- | --- | --- | --- | --- |
|  | 3 DPI | | 6 DPI | | 8 DPI | |
| Cytokine | Mock | IAV | Mock | IAV | Mock | IAV |
| IFNα | 434.14 ± 69.37 | 449.02 ± 58.83 | 1221.77 ± 590.9 | 2153.13 ± 707.64 | 332.16 ± 32.41 | 370.25 ± 17.28 |
| IFNγ | 8.15 ± 1.24 | 8.89 ± 1.12 | 19.75 ± 6.06 | 326.1 ± 99.35 | 3.89 ± 0.52 | 5.19 ± 1.17 |
| IL-28 | 14845.58 ± 2505.48 | 13601.43 ± 1964.5 | 8828.53 ± 2244.8 | 9710.21 ± 2314.45 | 13721 ± 1588.91 | 17267.6 ± 478.63 |
| IL-1α | 606.42 ± 136.54 | 502.4 ± 97.88 | **124.76 ± 32.02** | **434.1 ± 29.5** | 158.61 ± 23.99 | 155.23 ± 22.64 |
| IL-1β | 66.96 ± 8.74 | 44.84 ± 7.74 | **91.39 ± 15.48** | **499.69 ± 48.97** | 22.88 ± 3.25 | 21.89 ± 3.21 |
| IL-18 | **769.05 ± 97.94** | **326.1 ± 40.94** | **451.46 ± 125.05** | **3651.4 ± 300.64** | 229.95 ± 32.08 | 289.05 ± 16.12 |
| IL-33 | 9964.32 ± 2073.52 | 12244.06 ± 1311.09 | 16154.48 ± 4137.5 | 18531.12 ± 3165.64 | 6154.56 ± 1529.29 | 8396.15 ± 1579.39 |
| IL-33R | 111.27 ± 25.14 | 103.84 ± 25.44 | **100.98 ± 22.98** | **7827.65 ± 273.98** | 1547.53 ± 161.65 | 2357.48 ± 229.02 |
| IL-2 | 183.8 ± 24.37 | 167.56 ± 14.49 | **447.86 ± 85.84** | **959.35 ± 30.37** | 180.05 ± 8.07 | 156.58 ± 4.25 |
| IL-2R | 31.69 ± 3.65 | 32.24 ± 3.4 | 56.54 ± 4.2 | 60.11 ± 3.76 | 6.41 ± 0.6 | 6.94 ± 0.51 |
| IL-4 | 175.91 ± 16.41 | 174.3 ± 19.13 | 142.13 ± 26.51 | 65.78 ± 17.71 | 77.19 ± 6.58 | 82.45 ± 5.24 |
| IL-7 | 292.5 ± 63.45 | 249.14 ± 45.52 | 1014.69 ± 204.78 | 430.98 ± 158.57 | 213.08 ± 12 | 218.16 ± 12.15 |
| IL-7Rα | 884.75 ± 153.64 | 613.84 ± 118.3 | 1829.16 ± 356.25 | 1929.14 ± 391.14 | 567.12 ± 139.11 | 547.52 ± 114.2 |
| IL-9 | 274.36 ± 69.65 | 200.01 ± 39.27 | **182.17 ± 23.85** | **79.38 ± 5.16** | 147.31 ± 30.7 | 105.17 ± 21.14 |
| IL-15/ IL-15R | 41.19 ± 5.18 | 58.59 ± 19.14 | **7.84 ± 1.09** | **449.12 ± 106.44** | 41.14 ± 12.04 | 39.1 ± 9.55 |
| TNFα | 38.67 ± 7.83 | 34.08 ± 5.27 | **44.02 ± 4.97** | **340.84 ± 65.13** | 28.35 ± 6.44 | 32.25 ± 6.09 |
| RANKL | 48.87 ± 14.91 | 43.15 ± 4.07 | 30.34 ± 12.51 | 38.86 ± 15.78 | 19.15 ± 2.05 | 24.63 ± 1.8 |
| BAFF | 149.71 ± 25.58 | 154.74 ± 24.8 | 353.85 ± 108.43 | 336.08 ± 85.79 | 55.38 ± 10.02 | 55.83 ± 13.48 |
| IL-10 | 536.14 ± 152.25 | 507.22 ± 83.63 | 413.48 ± 226.16 | 479.6 ± 200.6 | 48.84 ± 15.93 | 53.87 ± 16.28 |
| IL-19 | 6176.59 ± 490.48 | 5805.54 ± 550.72 | 14908.86 ± 2400.6 | 15045.19 ± 2168.6 | 1636.23 ± 73.49 | 1928.8 ± 171.1 |
| IL-22 | 478.2 ± 75.22 | 330.12 ± 67.12 | **353.1 ± 77.97** | **1241.43 ± 169.88** | 265.48 ± 68.92 | 175.37 ± 38.56 |
| IL-12p70 | 18.07 ± 2.41 | 16.18 ± 1.11 | **17.61 ± 1.93** | **49.94 ± 5.74** | 12.78 ± 1.22 | 12.19 ± 1.12 |
| IL-23 | 2901.16 ± 278.62 | 2889.69 ± 228.03 | 2969.61 ± 614.09 | 2786.04 ± 618.41 | 2277.05 ± 160.36 | 2060.81 ± 205.15 |
| IL-27 | 160.24 ± 44.03 | 184.3 ± 39.6 | 324.32 ± 52.36 | 340.7 ± 50.87 | 47.55 ± 14.7 | 61.17 ± 13.26 |
| IL-17A | 173.79 ± 51.09 | 124.96 ± 34.21 | **219.69 ± 38.24** | **700.6 ± 55.14** | 256.74 ± 40.16 | 248.34 ± 28.7 |
| IL-25 | 342.51 ± 90.51 | 400.3 ± 73.64 | 585.14 ± 184.18 | 528.73 ± 143.7 | 111.03 ± 30.4 | 114.72 ± 26.24 |
| IL-6 | 154.51 ± 26.06 | 166.34 ± 21.96 | **26.83 ± 5.02** | **2176.7 ± 384.28** | 982.62 ± 133.95 | 1430.98 ± 98.25 |
| IL-31 | 1047.82 ± 256.71 | 1154.09 ± 155.5 | 1588.99 ± 337.71 | 1832.68 ± 288.98 | 1654.63 ± 71.97 | 1603.23 ± 83.81 |
| LIF | 70.88 ± 11.54 | 80.41 ± 8.7 | 117.28 ± 3.95 | 123.29 ± 3.87 | 117.99 ± 12.58 | 122.94 ± 15.56 |
| CCL2 | 970.74 ± 104.45 | 628.77 ± 53.68 | **249.36 ± 71.76** | **829.66 ± 70.47** | 454.52 ± 128.55 | 637.2 ± 148.41 |
| CCL3 | 44.43 ± 8.28 | 57.46 ± 8.24 | 0.81 ± 0.11 | 2.46 ± 0.52 | 10.62 ± 2.03 | 13.4 ± 2.59 |
| CCL4 | 36.44 ± 8.83 | 34.73 ± 6.37 | 40.93 ± 6.71 | 61.93 ± 10.78 | 15.24 ± 2.84 | 13.67 ± 1.75 |
| CCL5 | **8.96 ± 1.39** | **199.69 ± 44.62** | 3.99 ± 0.78 | 3.87 ± 0.72 | 2.54 ± 0.34 | 2.31 ± 0.34 |
| CCL7 | 401.91 ± 26.54 | 445.31 ± 28.89 | 353.1 ± 60.64 | 584.94 ± 34.76 | 228.49 ± 10.21 | 260.29 ± 8.12 |
| CCL11 | 298.65 ± 47.11 | 486.77 ± 62.86 | 521.57 ± 219.34 | 450.7 ± 71.84 | 248.62 ± 17.45 | 273.01 ± 14.86 |
| CXCL1 | 350.62 ± 38.53 | 239.36 ± 24.57 | 108.62 ± 15.04 | 100.32 ± 17.74 | 61.53 ± 9.25 | 82.3 ± 8.59 |
| CXCL2 | 57.11 ± 9.18 | 45.38 ± 5.52 | 36.43 ± 7.82 | 42.41 ± 7.28 | 16.74 ± 2.56 | 21.86 ± 5.22 |
| CXCL5 | 1355.95 ± 289.17 | 1581.24 ± 200.71 | 2672.42 ± 663.41 | 2621.18 ± 380.46 | 860.74 ± 90.65 | 854.6 ± 89.85 |
| CXCL10 | 187.75 ± 27.17 | 205.32 ± 24.39 | 352.68 ± 94.85 | 661.2 ± 94.48 | 178.65 ± 18.08 | 153.02 ± 17.45 |
| BTC | 278.46 ± 56.02 | 289.8 ± 49.28 | 222.58 ± 101.58 | 220.47 ± 73.35 | 49.01 ± 5.84 | 40 ± 8.48 |
| G-CSF | 12.98 ± 1.9 | 13.92 ± 1.88 | 136.68 ± 52.93 | 196.62 ± 54.92 | **24.75 ± 2.37** | **115.57 ± 16.54** |
| GM-CSF | **134.99 ± 22.56** | **52.97 ± 10.6** | 14.56 ± 4.59 | 13.52 ± 4.07 | 56.07 ± 12.96 | 44.09 ± 0.71 |
| M-CSF | 78.17 ± 14.16 | 85.54 ± 16.07 | **24.06 ± 6.55** | **99.11 ± 10.76** | **8.49 ± 2.64** | **54.52 ± 7.75** |
| IL-3 | 14.57 ± 2.98 | 10.93 ± 1.92 | 23.45 ± 5.23 | 22.04 ± 4.19 | 23.25 ± 3.89 | 18.05 ± 2.97 |
| VEGF-A | 372.26 ± 86.01 | 329.87 ± 68.39 | 2531.77 ± 525.28 | 2390.35 ± 494.67 | 209.14 ± 48.03 | 298.69 ± 51.04 |
| IL-13 | **99.43 ± 11.47** | **30.91 ± 5.11** | 82.79 ± 34.94 | 94.74 ± 37.37 | 29.75 ± 5.45 | 29.5 ± 5.79 |
| IL-5 | **138.65 ± 28.56** | **108.8 ± 13.45** | **524.01 ± 72.28** | **80.05 ± 21.27** | 80.34 ± 15.66 | 88.77 ± 15.44 |
| Leptin | 155.49 ± 41.44 | 189.16 ± 63.27 | 152.95 ± 28.22 | 161.83 ± 39.5 | **68.32 ± 26.23** | **823.7 ± 154.08** |

Table S4.

Bold indicates a statistically significant difference between mock and IAV at a given DPI based on multiple unpaired t tests with Holm-Šídák correction for multiple comparisons, adjusted p value <0.05 (n=12-14)
