## Supplemental Table 5 for "Suppression of progesterone by influenza A virus mediates adverse maternal and fetal outcomes in mice"

|  | Concentration in placenta (pg/ml) | | | |
| --- | --- | --- | --- | --- |
| Cytokine | Mock – Vehicle | IAV – Vehicle | Mock – 17-OHPC | IAV – 17-OHPC |
| IFNα | 2953.56 ± 345.71 | 3080.14 ± 316.04 | 2291.55 ± 388.78 | 2299.32 ± 297.92 |
| IFNγ | 20.28 ± 3.86 | **322.01 ± 92.93*** | 26.74 ± 3.9 | **28.29 ± 4.8#** |
| IL-28 | 10522.43 ± 1671.83 | 7253.82 ± 1349.19 | 10289.92 ± 1941.75 | 11093.45 ± 1152.45 |
| IL-1α | **156.56 ± 25.7** | **456 ± 24.33*** | 124.11 ± 17.23 | **183.26 ± 20.9#** |
| IL-1β | **200.13 ± 18.52** | **394.32 ± 41.67*** | 198.39 ± 24.16 | **190.68 ± 23.55#** |
| IL-18 | **594.03 ± 138.51** | **3641.84 ± 468.38*** | 608.28 ± 129.37 | **557.36 ± 74.18#** |
| IL-33 | 16742.25 ± 2192.21 | 21330.51 ± 2028.17 | 21105.58 ± 3615.48 | 22532.99 ± 4408.2 |
| IL-33R | **141.69 ± 19.73** | **8218.39 ± 219.98*** | 118.31 ± 16.18 | **116.74 ± 21.86#** |
| IL-2 | **568.95 ± 83.94** | **965.84 ± 25.61*** | 513.17 ± 73.41 | **467.62 ± 54.11#** |
| IL-2R | 65.27 ± 2.83 | 69.69 ± 2.71 | 57.33 ± 3.9 | 62.35 ± 2.49 |
| IL-4 | 150.67 ± 18.05 | 90.76 ± 8.1 | 173.67 ± 25.08 | 165.98 ± 19.48 |
| IL-7 | **1278.7 ± 150.43** | **412.44 ± 55.02*** | 1129.45 ± 182.31 | **1313.49 ± 153.31#** |
| IL-7Rα | 2707.53 ± 163.33 | 2813.94 ± 194.3 | 2307.3 ± 235.44 | 2033.51 ± 264.15 |
| IL-9 | **194.51 ± 16.46** | **90.03 ± 2.54*** | 226.46 ± 13.55 | **204.03 ± 19.87#** |
| IL-15/ IL-15R | **8.82 ± 0.94** | **571.21 ± 95.36*** | 8 ± 0.88 | **8.7 ± 0.79** |
| TNFα | **46.25 ± 3.74** | **395.91 ± 40.94*** | 48.66 ± 3.17 | **58.71 ± 6.48#** |
| RANKL | 43.3 ± 7.85 | 70.19 ± 13.52 | 63.66 ± 6.14 | 50.18 ± 8.68 |
| BAFF | 504.53 ± 91.44 | 467.26 ± 44.21 | 577.66 ± 62.77 | 409.53 ± 102.63 |
| IL-10 | 933.21 ± 127.07 | 922 ± 145.12 | 1000.85 ± 118.08 | 1103.29 ± 120.89 |
| IL-19 | 18157.95 ± 2088.25 | 15639.03 ± 1249.41 | 19405.69 ± 1433.86 | 15990.23 ± 1902.76 |
| IL-22 | **368.55 ± 60.61** | **1410.97 ± 157.15*** | 388.5 ± 81.27 | **462.39 ± 59.33#** |
| IL-12p70 | **19.61 ± 1.79** | **58.72 ± 5.45*** | 18.71 ± 0.86 | **15.9 ± 1.39#** |
| IL-23 | **3039.9 ± 426.18** | **3383.95 ± 462.82*** | 4016.9 ± 434.26 | **3654.11 ± 621.8#** |
| IL-27 | **429.58 ± 46.06** | **414.46 ± 43.66*** | 375.45 ± 31.54 | **397.81 ± 46.34#** |
| IL-17A | **215.19 ± 29.36** | **689.78 ± 24.77*** | 267.33 ± 22.13 | **210.56 ± 26.27#** |
| IL-25 | 592.51 ± 175.22 | 808.59 ± 144.54 | 833.26 ± 143.17 | 1045.33 ± 131.58 |
| IL-6 | **35.2 ± 5.68** | **1107.37 ± 233.81*** | 37.5 ± 6.58 | **38.38 ± 6.24#** |
| IL-31 | 1798.69 ± 272.53 | 2034.8 ± 180.84 | 1466.87 ± 251.8 | 1959.66 ± 159.88 |
| LIF | 124.96 ± 2.82 | 130.8 ± 3.55 | 126.62 ± 1.93 | 121.55 ± 3.84 |
| CCL2 | **337.34 ± 65.47** | **880.66 ± 63.71*** | 281.88 ± 72.71 | **345.65 ± 58.17#** |
| CCL3 | **0.91 ± 0.08** | **2.72 ± 0.43*** | 1.01 ± 0.07 | **1.13 ± 0.06#** |
| CCL4 | 52.44 ± 5.23 | 68.87 ± 8.41 | 56.63 ± 3.3 | 41.53 ± 5.4 |
| CCL5 | 3.12 ± 0.4 | 3.78 ± 0.48 | 4.99 ± 0.65 | 4.71 ± 0.61 |
| CCL7 | **343.12 ± 50.73** | **677.47 ± 16.7*** | 389.5 ± 46.45 | **479.78 ± 32.43#** |
| CCL11 | 1039.16 ± 154.72 | 528.98 ± 58.08 | 725.19 ± 183.71 | 904.7 ± 130.18 |
| CXCL1 | 129.04 ± 11.83 | 145.96 ± 7.46 | 131.42 ± 13.86 | 129.7 ± 10.67 |
| CXCL2 | 50.75 ± 6.04 | 58.94 ± 6.2 | 49.06 ± 7.41 | 40.8 ± 5.9 |
| CXCL5 | 4006.64 ± 257.2 | 2736.96 ± 346.74 | 4036.46 ± 370.3 | 2585.79 ± 617.36 |
| CXCL10 | 511.97 ± 68.28 | 739.96 ± 77.7 | 475.41 ± 73.49 | 460.24 ± 86.43 |
| BTC | 403.85 ± 61.18 | 265.84 ± 50.85 | 379.92 ± 56.81 | 320.84 ± 66.91 |
| G-CSF | 212.89 ± 48.31 | 281.79 ± 39.96 | 255.85 ± 32.76 | 244.71 ± 46.91 |
| GM-CSF | 22.51 ± 3.48 | 23.27 ± 2.59 | 17.37 ± 3.66 | 16.15 ± 3.84 |
| M-CSF | **31.92 ± 4.15** | **103.25 ± 6.75*** | 25.36 ± 6.24 | **26.32 ± 5.22#** |
| IL-3 | 29.04 ± 3.95 | 24.24 ± 3.19 | 28.87 ± 4.25 | 25.59 ± 3.55 |
| VEGF-A | 3026.72 ± 378.13 | 2966.6 ± 371.77 | 2698.08 ± 457.09 | 3140.29 ± 426.41 |
| IL-13 | 126.35 ± 25.73 | 101.78 ± 26.56 | 111.33 ± 22.27 | 126.24 ± 23.27 |
| IL-5 | **659.99 ± 46.8** | **95.87 ± 18.37*** | 627.28 ± 48.55 | **660.35 ± 62.75#** |
| Leptin | 184.66 ± 20.55 | 202.79 ± 30.19 | 189.82 ± 25.96 | 187.46 ± 17.51 |

Table S5.

Bold indicates a statistically significant differences, with significant differences between vehicle treated mock and IAV infected groups indicated with an asterisk (*) and significant differences between IAV infected vehicle and 17-OHPC treated groups indicated with a pound sign (#) based on two-way ANOVA with Bonferroni post hoc test, p value <0.05 (n=9/group)
